# Modularity of PRC1 Composition and Chromatin Interaction define Condensate Properties

**DOI:** 10.1101/2023.10.26.564217

**Authors:** Stefan Niekamp, Sharon K. Marr, Theresa A. Oei, Radhika Subramanian, Robert E. Kingston

## Abstract

Polycomb repressive complexes (PRC) play a key role in gene repression and are indispensable for proper development. Canonical PRC1 forms condensates *in vitro* and in cells and the ability of PRC1 to form condensates has been proposed to contribute to maintenance of repression. However, how chromatin and the various subunits of PRC1 contribute to condensation is largely unexplored. Using single-molecule imaging, we demonstrate that nucleosomal arrays and PRC1 act synergistically, reducing the critical concentration required for condensation by more than 20-fold. By reconstituting and imaging PRC1 with various subunit compositions, we find that the exact combination of PHC and CBX subunits determine the initiation, morphology, stability, and dynamics of condensates. In particular, the polymerization activity of PHC2 strongly influences condensate dynamics to promote formation of structures with distinct domains that adhere to each other but do not coalesce. Using live cell imaging, we confirmed that CBX properties are critical for condensate initiation and that PHC polymerization is important to maintain stable condensates. Together, we propose that PRC1 can fine-tune the degree and type of condensation by altering its composition which might offer important flexibility of regulatory function during different stages of development.

## Introduction

Chromatin structure and accessibility plays a critical role in determining gene expression programs that underlie cellular fates and functions in development and disease^1–4^. In eukaryotes DNA is wrapped around histone octamers and generally organized into two different types of nuclear domains: An accessible, “open” state that allows for transcription to occur and an inaccessible, “closed” state that is associated with gene repression^5,6^. To establish, maintain, or alter these chromatin states, eukaryotes employ chromatin associated proteins. One such family of proteins are the Polycomb group (PcG) proteins which are key players to maintain the repression of hundreds of genes and are essential for proper development^7–12^. PcG proteins are typically found in one of two major families of complexes that modify histones and chromatin structure: Polycomb repressive complex 1 (PRC1) and PRC2. In particular, PRC1 contributes to local and long-range chromatin compaction, resulting in cellular foci called Polycomb bodies^13–17^. The canonical form of PRC1 has been shown to form condensates *in vitro* and in cells and the ability of PRC1 to form condensates that concentrate components of the Polycomb-Group has been proposed to contribute to maintenance of repression^8–10,18,19^.

In recent years, formation of condensates via phase separation has emerged as an organizing principle for macromolecules in the nucleus^2,20–26^. Phase separation has been linked to transcription factors, RNA polymerase II, and coactivators that activate transcription^27–35^, as well as to heterochromatin protein 1 (HP1) alpha and canonical PRC1 that aid in transcriptional repression^18,19,36–38^. In addition to the condensate formation of chromatin modifying and binding proteins, chromatin itself has been shown to form condensates^22,24,25,39,40^. However, how PRC1 condensates are organized in the context of chromatin and whether their interaction is synergistic is unknown.

Canonical PRC1 consists of four proteins: RING, PCGF, CBX, and PHC. Of these four subunits, CBX and PHC are thought to be the main drivers that alter chromatin structure^7–12^. Each of these four subunits has multiple paralogs resulting in over 60 possible combinations for canonical PRC1 alone (**Figure S1A**). In addition, the various paralogs have a dynamic expression pattern in different cell types and change during differentiation^13,41^. Dissecting the contributions of various paralogs on chromatin compaction and condensation has been challenging in cells because of their redundancy. For instance, the chromobox (CBX) protein has five paralogs (CBX2, CBX4, CBX6, CBX7, and CBX8) of which some are known to drive Compaction and Phase Separation via the CaPS region^8–10,18,19,42^. Point mutations in the highly positively charged region of the CBX2 CaPS region impair its compaction and phase separation ability resulting in developmental phenotypes in mice^43–45^. In addition to the CBX subunit the PHC subunit has also been implicated in condensate formation^38^. The PHC subunit has a conserved sterile alpha motif (SAM) at its C-terminus that can form head-to-tail polymers^46^. Point mutations inhibiting the polymerization activity of PHC have been linked with decreased repression in mice and knock-out studies in cells have shown that PHC polymerization is essential to maintain long-range chromatin interactions that connect different regions of the genome^13–15^. However, the exact contributions and the molecular mechanism of each of the CBX and PHC subunits in a full PRC1 are poorly understood. It is unknown if either CBX and PHC have a dominant effect, if both have additive effects, or if they contribute different biochemical properties to achieve condensation.

Here, we used a reconstitution approach and showed that condensate formation and condensate properties depend on the specific combination of PRC1 subunits. Using single-molecule and live cell imaging, we found that the properties of the CBX and PHC subunits determine condensate initiation, morphology, stability, and dynamics. Specifically, the PHC subunit alters morphology and dynamics of PRC1 condensates which results in the formation of distinct and long-lived domains. The ability of PHC2 to drive formation of these altered condensates is disrupted by a point mutation in the SAM domain that previously was shown to lead to homeotic transformation in mice^14^. Moreover, we found that the CBX properties are critical for condensate initiation. Mutations in a basic patch of CBX2 that abolish condensate formation were also shown to lead to homeotic transformation in mice^19,44^. In addition, we defined the dynamics of condensate formation in real time with single-molecule resolution by using a newly developed TIRF microscopy-based approach. We found that nucleosomal arrays reduce the concentration at which PRC1 forms condensates by more than 20-fold. Furthermore, we observed that a sufficient nucleosomal array length and stoichiometric excess of PRC1 to nucleosomes are required for condensate formation. Together, the data suggests that the synergistic interaction of chromatin and PRC1 drives condensate formation and that PRC1 composition can fine-tune condensate properties. We propose that the range of PRC1 condensate properties is an important mechanism by which PRC1 determines the degree of gene repression and thereby regulates developmental processes.

## Results

### Polymerization activity of PHC2 changes condensate morphology and stability

To tease apart the contributions of different PRC1 subunits on condensate formation with nucleosomal arrays we focused on two families of subunits that have previously been shown to form condensates: CBX and PHC^18, 19,38^. We first focused on the PHC subunit and employed an epifluorescence microscopy-based assay with recombinant PRC1. In this assay we mixed fluorescently-labeled 12-mer nucleosomal arrays with the 5S-sequence^47,48^ with canonical PRC1 composed of RING1B, BMI1, mGFP-CBX2 and PHC2 (**Figures 1A, S1A, B**). We compared complexes formed with wild type PHC2 (PHC2 WT) to those formed with the PHC2 polymerization deficient mutant L307R (PHC2 L307R)^14,38,46^ (**Figures 1A, B, S1A-D**). The PHC2 L307R mutant was chosen because it has a homeotic phenotype in mice, indicating that it is a hypomorph for PcG function in development^14^. It has previously been used in functional analysis of PRC1 because the mutation prevents polymerization of PRC1 into insoluble aggregates^14,38,46^. To perform assays with PHC2 WT, we added a HALO-tag separated by a TEV site to the C-terminus of PHC2 WT and PHC2 L307R, which sterically blocks polymerization of PHC2 WT^14^. This enabled the purification of monomeric PRC1. Adding TEV protease to cleave the HALO-tag at the start of the experiment allowed polymerization of the PHC2 SAM domain^14^ (**Figures 1A, S1E**). Thus, the addition of a cleavable polymerization blocker allowed comparison of the role for WT and polymerization deficient PHC2 on condensate formation of intact PRC1.

**Figure 1.**
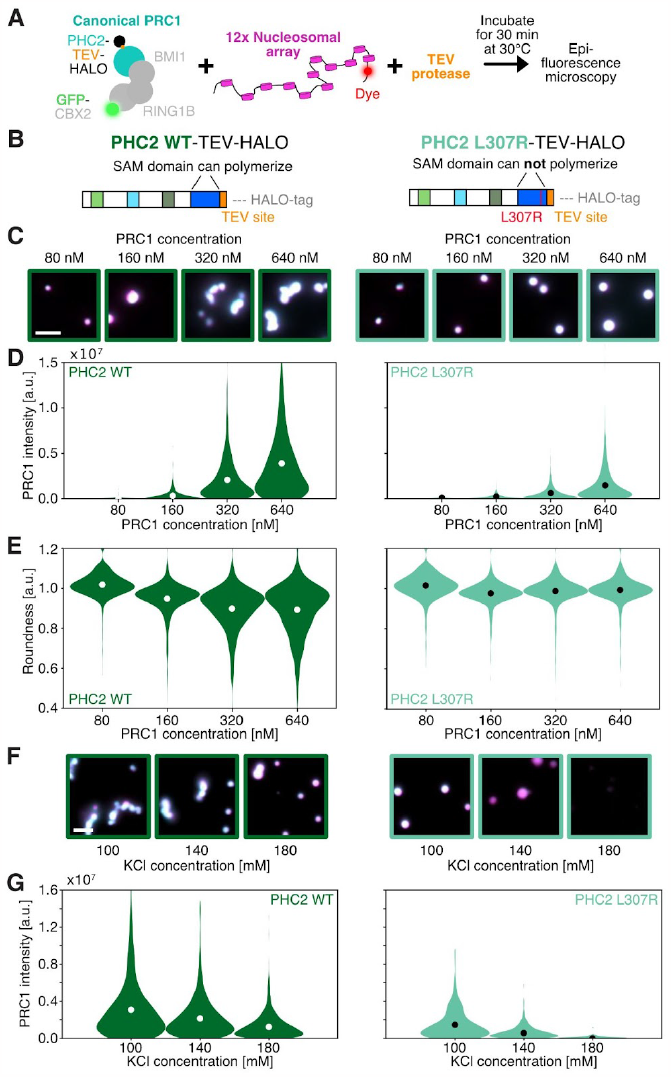
PHC polymerization alters condensate morphology and stability. (**A**) Schematics and work-flow of epifluorescence microscopy assay to study condensate formation of nucleosomal arrays and canonical PRC1 (for more details see **Figure S1A-E**). (**B**) Schematics of different PHC constructs. (**C**) Example micrographs of nucleosomal arrays (magenta) and canonical PRC1 complexes (cyan) with different subunits incorporated. The scale bar is 5 μm. (**D**) Violin plot of average PRC1 intensities for different PRC1 compositions as shown in **C**. The nucleosome concentration was 30 nM. Circles represent the average value. (**E**) Violin plots of roundness values for different PRC1 compositions. Circles represent the average value. (**F**) Example micrographs of nucleosomal arrays (magenta) and canonical PRC1 complex (cyan) with mGFP-tagged CBX2 and either PHC2 WT (dark green box) or PHC2 L307R (light green box) at different potassium chloride concentrations. Nucleosome concentration was 30 nM and PRC1 concentration was 610 nM. The scale bar is 5 μm. (**G**) Violin plot of PRC1 intensities of data as shown in **F**. Circles represent the mean average value. (**D, E, G**) For each condition 325 to >1000 spots have been analyzed. Data are representative of at least three technical repeats.

We first focused on comparing PHC2 WT and L307R in the background of the full PRC1 complex containing RING1B, BMI1, and CBX2 WT and subsequently tested the role of changing CBX subunits. When we mixed different PRC1 concentrations with nucleosomal arrays and imaged these with an epifluorescence microscope after a 30-minute incubation at 30°C we observed a striking morphology difference for PRC1 complexes with PHC2 WT compared to the PHC2 L307R mutant (**Figure 1C**). Condensates of a PRC1 complex with PHC2 L307R appeared as almost perfectly round spheres while a PRC1 complex with PHC2 WT often had a long, sometimes branched shape (**Figure 1C-E**). We quantitatively analyzed the roundness of these condensates (**Figure S1F**) and found that the average roundness for a PRC1 complex with PHC2 L307R is 0.99 (equivalent to a perfect circle), while the average roundness of PRC1 complexes containing PHC2 WT measured 0.88 (closer to a square than a circle) (**Figure 1E**). Together, we conclude that the polymerization activity of PHC2 alters the morphology of condensates formed by PRC1 and nucleosomal arrays.

Since the ionic strength of buffers has been shown to influence condensate formation of nucleosomal arrays and PRC1^19,25^, we asked whether different salt concentrations could result in different outcomes for condensate formation of polymerization competent versus deficient PHC2. To test this, we mixed PRC1 complexes containing either PHC2 WT or PHC2 L307R with nucleosome arrays in buffers with 100, 140, or 180 mM potassium chloride. We then performed the same imaging assay as described above. We found that the intensity of the PHC2 L307R mutant-based condensates decreased with higher salt concentrations and observed almost no condensates at 180mM KCl (**Figure 1F**). In contrast, nucleosomal array and PRC1 intensity remained almost constant for PHC2 WT over all salt concentrations (**Figures 1G, S1G**). Moreover, we found that the roundness for PHC2 WT condensates increased with higher salt concentrations, and that PHC2 WT maintained a similar fold difference in roundness compared to PHC2 L307R mutant PRC1 over all salt concentrations (**Figure S1H**). Thus, we conclude that the polymerization activity of PHC2 can increase the stability of condensates at physiological salt concentrations.

### PHC2 polymerization activity restricts dynamics of PRC1 condensates

To assess whether PHC polymerization activity alters the dynamics of condensates, we developed a three-color imaging assay to determine if PRC1 molecules and nucleosomal arrays can exchange between condensates. We first focused on the exchange of PRC1 molecules and asked if PRC1 molecules in condensates of one color can exchange with PRC1 molecules in condensates of different color (**Figure 2A**). Specifically, we used mGFP-tagged PRC1 complexes (cyan) and PRC1 complexes that were TMR labeled via HALO-tag^49^ (yellow). Both differently colored PRC1 complexes were prepared separately using nucleosomal arrays in which DNA was labeled with an ATTO 647N dye (red). After a sufficiently long incubation (30 minutes) for condensate formation to reach a steady state, we had a preparation of condensates with red nucleosomal arrays and cyan PRC1 complexes and a separate preparation of condensates consisting of red nucleosomal arrays and yellow PRC1 complexes. We then mixed these two preparations together and incubated them for 1, 15, 30, and 60 minutes (**Figure 2A**). If the PRC1 molecules exchanged, we would have expected to see the red nucleosomal arrays together with the cyan and yellow PRC1 complexes in the same condensate. However, we found that PRC1 complexes with PHC2 WT formed larger condensate clusters that maintained well-defined boundaries for the two differently colored PRC1 preparations, indicating that very limited mixing had occurred (**Figures 2B, S2A**). These condensate clusters remained distinct for the experimental duration of one hour (**Figures 2B, E, S2C**). This is in stark contrast to the polymerization deficient PHC2 L307R mutant where, after performing the analogous experiment, we observed high levels of mixing as demonstrated by the formation of PRC1 condensates containing both colors of complexes (**Figures 2B, E, S2B, D**).

**Figure 2.**
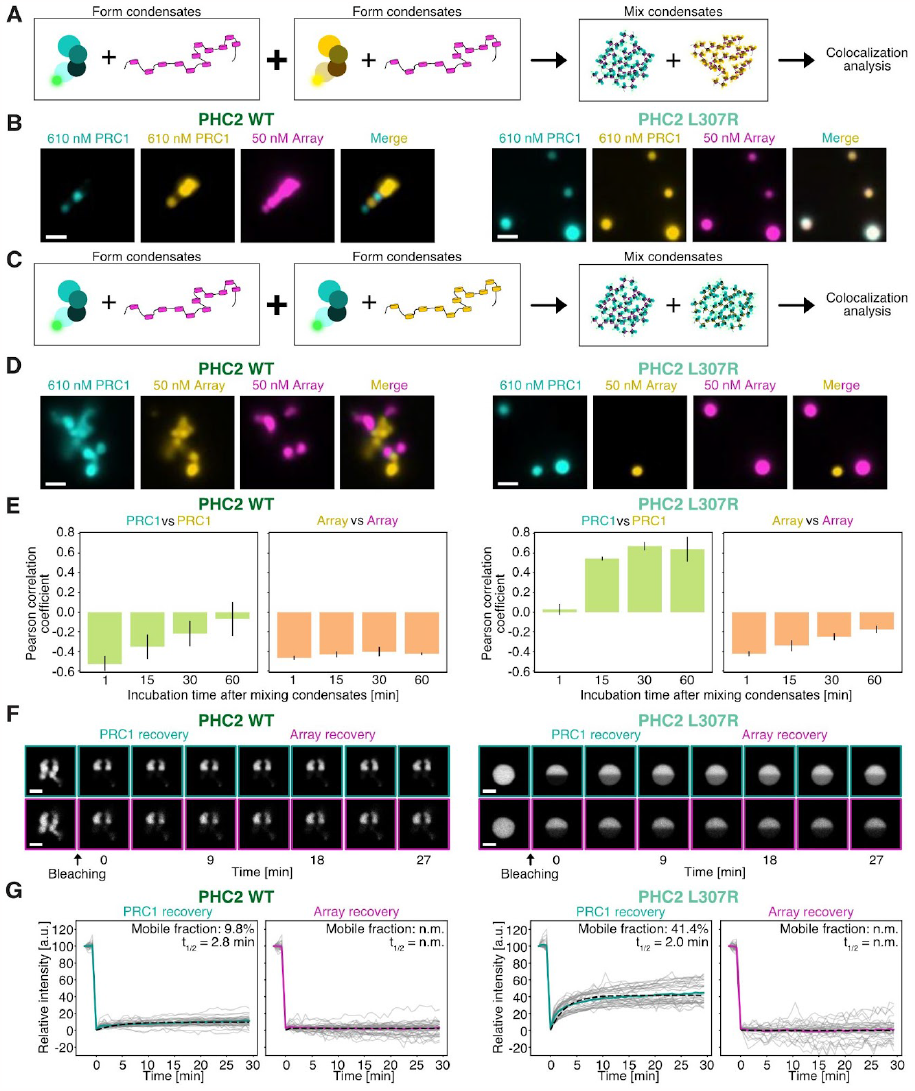
SAM polymerization activity of PHC2 alters PRC1 dynamics. (**A**) Schematic for mixing experiments of the full canonical PRC1 complex with Alexa 488 (cyan) or TMR (yellow) labeled HALO-tag of the CBX2 subunit. (**B**) Representative micrographs with differently colored PRC1 molecules 30 minutes after mixing for PRC1 complexes with PHC2 WT and PHC2 L307R. The scale bar is 5 μm. (**C**) Schematic for mixing experiments of ATTO-647N (magenta) or Cy3N (yellow) labeled nucleosomal arrays. Here, CBX2 of PRC1 was mGFP-tagged and the PHC2 HALO tag was unlabeled. (**D**) Representative micrographs of condensates with differently colored nucleosomal arrays 30 minutes after mixing for PRC1 complexes with PHC2 WT and PHC2 L307R. The scale bar is 5 μm. (**E**) Average Pearson correlation coefficients of differently colored PRC1 molecules (left) and nucleosomal arrays (right) for PRC1 complexes with PHC2 WT and PHC2 L307R. For each condition >100 spots have been analyzed. The error bar is the standard deviation of three repeats. (**F**) Images of nucleosomal arrays (magenta box) and canonical PRC1 (cyan box) with PHC2 WT and PHC2 L307R before and after bleaching. The scale bar is 2 μm. (**G**) FRAP trajectories for PRC1 (cyan) with PHC2 WT and PHC2 L307R and nucleosomal arrays (magenta). Colored trajectories are the average of all individual traces shown in gray. (**E, G**) Nucleosome concentration was 30 nM and PRC1 concentration was 610 nM. Data are representative of at least three technical repeats.

We quantitatively confirmed this mixing by measuring the colocalization between the two differently colored PRC1 complexes in condensates (Pearson correlation coefficient after one hour of mixing: PHC2 WT = −0.21; PHC2 L307R 0.74) (**Figures 2E, S2C, D**). Together, we conclude that the PHC2 polymerization activity limits the exchange of PRC1 molecules among condensates.

We next asked whether the exchange dynamics of nucleosomal arrays was also altered by the PHC2 polymerization activity. We employed a similar three-color imaging system, this time using mGFP-tagged PRC1 (cyan) and two colors of nucleosomal arrays in which DNA was either labeled with a Cy3N dye (yellow) or an ATTO 647N dye (red) (**Figure 2C**). Each array was separately combined with mGFP-PRC1 and allowed to form condensates. The differently colored condensates were then mixed and imaged. We observed that condensates formed with PHC2 WT containing PRC1 adhered to each other but did not fuse, giving rise to condensate clusters with well-defined boundaries (**Figures 2D, E, S2E, G**). This shows that very limiting mixing of nucleosomal arrays among condensates had occurred. We also saw very limited mixing for condensates with PRC1 complexes containing PHC2 L307R (**Figures 2D, E, S2F, H**). However, for the PHC2 L307R mutant the condensates with the differently colored arrays did not adhere to each other. We conclude that nucleosomal arrays do not exchange among condensates and that this is independent of the PHC2 polymerization activity. However, the PHC2 polymerization activity is required to bring different condensates of PRC1 and nucleosomal arrays together into larger clusters in which condensates stay in individual compartments.

The differences in mixing of the polymerization active and deficient PHC2 complexes suggest that the presence of an active polymerization domain in PHC2 might alter the physical properties of condensates from a liquid-like to a more gel-like state. To test this, we used Fluorescent Recovery After Photobleaching (FRAP) on condensates of nucleosomal arrays and PRC1 with either PHC2 WT or PHC2 L307R. In both cases the nucleosomal array intensity did not recover at all after bleaching (**Figure 2G**), supporting our observation that nucleosomal arrays in condensates are not dynamic. When we bleached PRC1 in condensates with PHC2 WT, we found that the PRC1 intensity recovered slowly and only to 9.8% of the pre-bleach intensity (**Figures 2F, G, S7I; Movies S1, 3**). In contrast, the PRC1 intensity of the PHC2 SAM mutant recovered to 41.1% of the pre-bleach intensity (**Figures 2F, G, S7J, Movies S2, 4**). Taken together, these data support the hypothesis that the polymerization activity of PHC2 reduces the mobility of PRC1 molecules and that the PHC polymerization shifts the condensates to a more gel-like state than a liquid-like state^23^. This transition might explain the changes in morphology, stability, and dynamics that we observed between PRC1 complexes with polymerization active and deficient PHC2.

### Properties of the CBX subunit are important for condensate formation

The CBX family of proteins is the other component of canonical PRC1 complexes that is known to contribute to condensate formation. To characterize that family, we focused on CBX2 and CBX7, as CBX2 displays strong compaction and condensate formation while CBX7 lacks these abilities at the same concentration^19,50^ (**Figure 3A, B**). To characterize the function of the CBX2 compaction and phase separation (CaPS) region we used a gain of function CBX7 chimera (CBX7 CaPS) that has been shown to form condensates^50^. We performed all experiments with the different CBX subunits in the background of PHC2 WT and PHC2 L307R and imaged condensate formation for all six PRC1 complex compositions of the CBX and PHC proteins being examined (**Figures 3C, S3A**).

**Figure 3.**
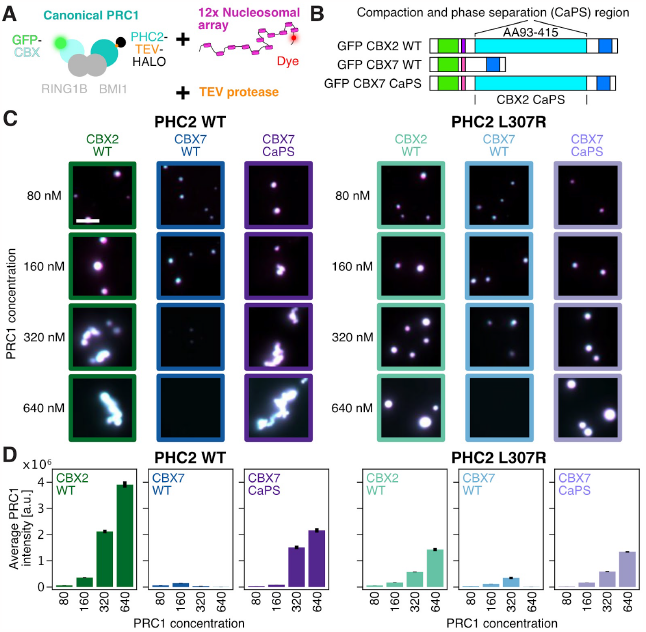
Contributions of CBX subunit to condensate formation. (**A**) PRC1 constructs with different PHC and CBX subunits are mixed with nucleosomal arrays and TEV protease. (**B**) Schematics of different CBX constructs. (**C**) Example micrographs of nucleosomal arrays (magenta) and canonical PRC1 complexes (cyan) with different subunits incorporated. The scale bar is 5 μm. (**D**) Average PRC1 intensities for different PRC1 compositions as shown in **C**. The nucleosome concentration was 30 nM. Error bar is standard error of the mean. For each condition 325 to >1000 spots have been analyzed. Data are representative of at least three technical repeats. The intensity data for PRC1 with CBX2 WT and PHC2WT or PHC2 L307R is the same as shown in **Figure 1D**.

When we compared the CBX subunits, we found that PRC1 complexes based on CBX7 WT either did not form condensates or formed only small condensates compared to CBX2 WT and CBX7 CaPS (**Figures 3C, D, S3B**). Even the brightest CBX7 WT condensates were only as bright as the dimmest condensates formed by CBX2 WT PRC1 containing complexes over the full PRC1 concentration regime. For instance, at 640 nM PRC1 concentration, we found a ∼5,000-fold increase in the intensity of condensates for CBX2 WT compared to CBX7 WT when both are in a complex with PHC2 WT, BMI1, and RING1B. For PRC1 complexes containing CBX7 with the CBX2 CaPS region inserted, we observed very similar condensate properties in terms of morphology and salt stability as for CBX2 WT containing PRC1 complexes (**Figures 3C, D, S3C-F**). Interestingly, the intensity of condensates with CBX7 WT did not peak at the highest PRC1 concentration as it did for CBX7 CaPS and CBX2 WT containing condensates (**Figure 3D**). This suggests that additional CBX7 complexes have a negative effect on condensate formation above a threshold concentration. One possible explanation might be that CBX7 coats nucleosomal arrays and prevents interactions among arrays at high CBX7 concentrations. Overall, we found that the ability to form condensates was independent of the PHC2 polymerization capability suggesting the CaPS region of the CBX is required to drive condensate formation in PRC1 complexes that contain PHC2 (**Figure 3C, D**). To test the requirement of CBX properties for condensate formation further, we used two additional, previously described loss of function mutants of the CBX2 CaPS region that have been shown to reduce chromatin compaction and PRC1 condensation^19,43,44^ (**Figure S3G**). In agreement with previous studies^19^, we found that the number of lysines and arginines in the CaPS region correlates with the PRC1 concentration at which condensates begin to emerge (**Figure S3H, I**). Taken together, the data suggests that the properties of the CBX subunit determine if condensates form in a PHC2-polymerization activity independent manner.

### PHC1 can create condensates independent of CBX properties

Since we observed striking differences in terms of condensate formation for the PHC polymerization activity and for the different CBX paralogs, we tested whether the PHC paralogs PHC1 and PHC2 would also show distinct behavior (**Figure 4A**). This is particularly interesting because CBX7 and PHC1 are more commonly expressed together in embryonic stem cells, which require developmental plasticity, while PRC1 complexes composed of CBX2 and PHC2 are more commonly found in differentiated cells^13,41^. Since embryonic stem cells have been shown to contain PRC1 condensates which contain CBX7, we analyzed whether PHC1 might contribute to condensate formation when paired with CBX7. We expressed PHC1 wild type (PHC1 WT) and a polymerization deficient version of PHC1 (PHC1 L994R), and purified all possible combinations of PHC1, PHC2, CBX2, and CBX7 to form full canonical PRC1 complexes with RING1B and BMI1. As with PHC2, the polymerization of PHC1 was initiated by cleaving off the C-terminal HALO-tag after purification of the complex and during mixing of PRC1 with nucleosomal arrays. We imaged the resultant samples using epifluorescence microscopy.

**Figure 4.**
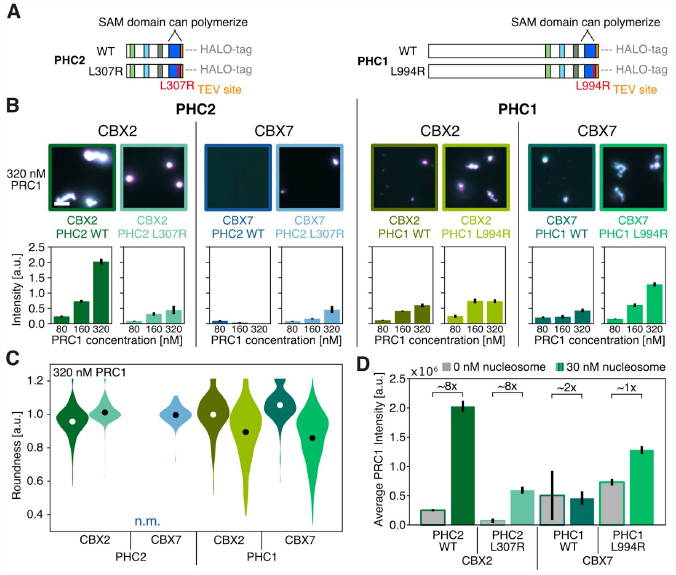
PRC1 complexes with PHC1 form condensates independent of CBX properties. (**A**) Schematics of different PHC constructs. (**B**) Top: Example micrographs of 30 nM nucleosomal arrays (magenta) and 320 nM canonical PRC1 complexes (cyan) with different subunits incorporated. The scale bar is 5 μm. Bottom: Average PRC1 intensities for different PRC1 compositions. The nucleosome concentration was 30 nM. Error bar is standard error of the mean. For each condition 45 to >750 spots have been analyzed. Data are representative of at least two technical repeats. (**C**) Violin plots of roundness values for different PRC1 compositions as shown in **B**. The nucleosome concentration was 30 nM. Circles represent the mean average value. (**D**) Average PRC1 intensities for different PRC1 compositions mixed with (different shades of green) and without (gray) nucleosomal arrays. The nucleosome concentration was 30 nM and the PRC1 concentration was 320 nM. Error bar is standard error of the mean. (**C, D**) For each condition 90 to >750 spots have been analyzed. Data are representative of at least three technical repeats. N.m. is not measurable.

We found that condensates containing PRC1 complexes with PHC1 looked strikingly different compared to PHC2 containing condensates (**Figures 4B**). For instance, we observed large condensates for PRC1 complexes containing CBX7 and PHC1, in contrast to the small condensates of PRC1 complexes containing CBX7 and PHC2. The PRC1 intensity at 160 nM for PRC1 complexes containing CBX7 and PHC1 was 10-fold higher than for complexes containing CBX7 and PHC2 suggesting that the PHC1 condensation properties can overcome the lack of CBX condensation abilities (**Figure 4B**). Moreover, the PRC1 intensities of complexes containing CBX2 and PHC1 were ∼4-fold lower than the PRC1 intensities of complexes containing CBX2 and PHC2, suggesting that the nature of the PHC subunit can also modulate condensate formation of PRC1 complexes containing CBX2 (**Figure 4B**). As for PHC2, we observed differences for the polymerization active and deficient versions of PHC1. However, this time PHC1 WT formed round condensates, while PHC1 L994R formed clusters of condensates that adhered to each other but did not coalesce (**Figure S4A, B**). Furthermore, we found that PHC1 creates more stable condensates with respect to the potassium chloride concentration than PHC2 does (**Figure S4C, D**). We conclude that PHC1 operates differently than PHC2 in terms of condensate formation, morphology, and dynamics. Furthermore, the data implies that PHC functionality is, in part, contingent on the specific CBX subunit with which they are associated.

Since PHC1 has a much longer N-terminus than PHC2 (**Figure 4A**), we wondered whether the different condensate properties among these two PHC paralogs can be attributed to the different N-termini. To test this, we removed the N-terminus of PHC1 and created a truncated PHC1 which has a similar size compared to PHC2 (**Figure S4E**). Imaging PRC1 complexes containing CBX7 and either full length or truncated PHC1 revealed that only complexes with the full length PHC1 can form condensates (**Figure S4F, G**). This data suggests that the N-terminus of PHC1 plays a role in the differences in condensate properties between PHC1 and PHC2, similar to the earlier studies with Drosophila PH^51^. We further conclude that PHC1 and PHC2 have significantly different condensation formation properties *in vitro* and that the extended N-terminus of PHC1 is involved in condensate formation. Interestingly, a previous study has shown that the N-terminus of PHC1 can be modified by the glycosyltransferase Ogt by adding O-linkedN-Acetyl-glucosamine (O-GlcNAc) which has been suggested to be essential for ordered PRC1 assemblies^52^. This modulation of PHC1 is not fully understood, so we chose to focus on PHC2 for the remaining experiments.

### Polymerization of PHC is required for condensate stability in cells

We tested our *in vitro* derived rules of PRC1 condensate formation in cells. To this end, we used NIH-3T3 fibroblast cell lines that expressed doxycycline-inducible mGFP-tagged CBX2 WT or mGFP-tagged CBX7 WT. In addition, we transiently expressed mScarlet-tagged PHC2 WT or mScarlet-tagged PHC2 L307R and performed confocal microscopy. For cells expressing mGFP-CBX2 WT and mScarlet-PHC2 WT we observed colocalization in condensates suggesting that CBX2 WT and PHC2 WT form condensates together (**Figure 5A-C**). In agreement with the *in vitro* data, we did not observe any condensates in cells expressing mGFP-CBX7 WT but only in cells expressing mGFP-CBX2 WT (**Figure 5C-F**). Strikingly, when we imaged cells co-expressing mGFP-CBX2 WT and mScarlet-PHC2 L307R we detected fewer CBX2 condensates than for cells co-expressing mScarlet-PHC2 WT (**Figure 5C, D**), consistent with what was observed in a human cell line^14^. Moreover, in neighboring cells that only express mGFP-CBX2 WT we could clearly observe many condensates which were likely formed with PRC1 complexes containing endogenous PHC protein. Since the PHC2 L307R can still form dimers but not long polymers^14^, it is likely that the mutant PHC2 L307R, when overexpressed, formed dimers with almost all endogenous PHC and prevented the endogenous PHC from polymerizing and thus stabilizing condensates. Measuring the number of condensates per nucleus revealed a ∼4-fold decrease for cells co-expressing mGFP-CBX2 WT and mScarlet-PHC2 L307R when compared to cells only expressing mGFP-CBX2 WT or mGFP-CBX2 WT together with mScarlet-PHC2 WT (**Figure 5G**). When condensates formed, they all had a similar average intensity irrespective of whether PHC2 was WT or mutant. This suggests that PHC2 polymerization activity does not impact condensate size but stability (**Figure 5H**). Together, the live cell imaging data is in good agreement with the *in vitro* observations presented above that condensate initiation depends on CBX properties and that PHC2 polymerization is required for condensate stability.

**Figure 5.**
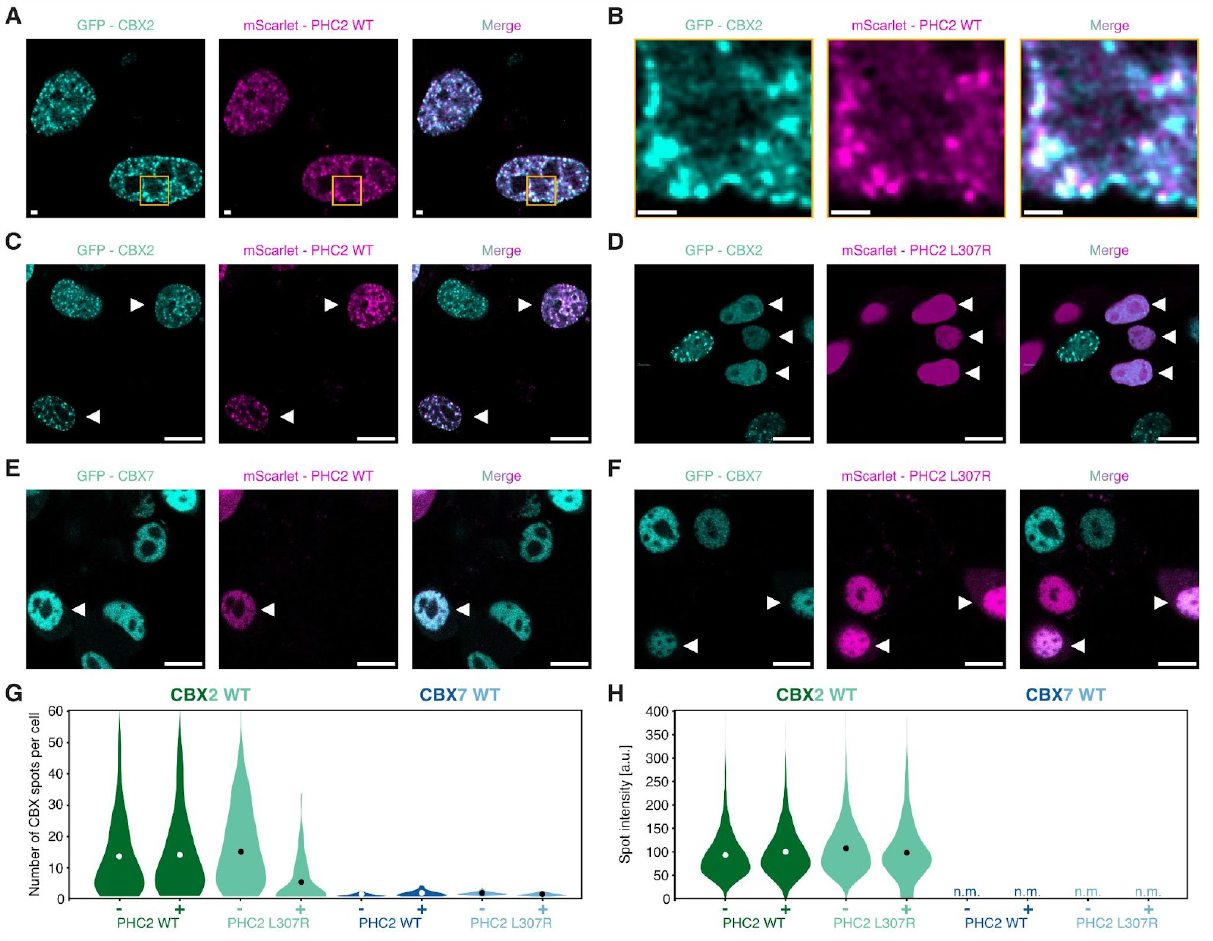
SAM polymerization of PHC2 is required for condensate stability in cells. (**A, B**) Deconvoluted confocal microscopy images of fibroblast 3T3 cells with dox induced expression of mGFP-CBX2 and transiently transfected mScarlet-PHC2 WT. Orange boxes on the left show areas that are magnified in **B**. Scale bar is 1 μm. (**C-F**) Deconvoluted confocal microscopy images of fibroblast 3T3 cells with dox induced expression of (**C, D**) mGFP-CBX2 or (**E, F**) mGFP-CBX7 and transiently transfected (**C, E**) mScarlet-PHC2 WT or (**D, F**) mScarlet-PHC2 L307R. White triangles indicate cells that have both CBX and PHC2 expression. Scale bar is 10 μm. (**G**) Violin plots with quantification of the number of spots per nucleus for cells expressing mGFP-CBX2 or mGFP-CBX7 and either with co-expression of mScarlet PHC2 (indicated by “+”) or not (indicated by “-”). Circles represent the mean average value. Data is pooled from 3 repeats and more than 50 cells were analyzed for each condition. (**H**) Violin plots with quantification of the intensities of spots for cells expressing mGFP-CBX2 or mGFP-CBX7 and either with co-expression of mScarlet PHC2 (indicated by “+”) or not (indicated by “-”). Circles represent the mean average value. Data is pooled from 3 repeats and more than 50 cells were analyzed for each condition. N.m. is not measurable.

### Nucleosomal arrays and PRC1 act synergistically to form condensates

In the nucleus PRC1 normally is bound to the nucleosomal arrays that form chromatin. Nucleosomal arrays are known to be able to form condensates in isolation^19^. We therefore characterized the impact of arrays on the ability of PRC1 to form condensates. To test this, we developed a single-molecule Total Internal Reflection Fluorescence (TIRF) microscopy protocol using a mobile, biotinylated lipid bilayer to visualize the dynamics of condensate formation in real time (**Figure 6A**). We tethered biotinylated DNA of fluorescently-labeled 12-mer nucleosomal arrays to biotinylated lipids via streptavidin, and allowed PRC1 to bind to nucleosomal arrays (**Figure 6B**). These tethered nucleosomal arrays can freely move in two dimensions above the bilayer. For control experiments with PRC1 alone and PRC1 with non-biotinylated nucleosomal arrays, we attached StrepII-tagged PRC1 to biotinylated lipids (**Figure 6B, Materials and Methods**). For our TIRF assay experiments we focused on PRC1 complexes containing PHC2 WT or PHC2 L307R together with mGFP-CBX2 WT, BMI1, and RING1B as these PRC1 compositions drive condensate formation (**Figure 3**). Together, these protocols allowed the detection of single nucleosomal arrays and condensates, enabled us to estimate the number of PRC1 molecules bound to nucleosomal arrays and in condensates, and facilitated direct imaging of condensate formation (**Figure S5A**). The single-molecule resolution of this assay allows visualization of condensates with size and intensity below the limit of regular epifluorescence microscopy setups and broadens analysis to more than 10-fold lower concentrations.

**Figure 6.**
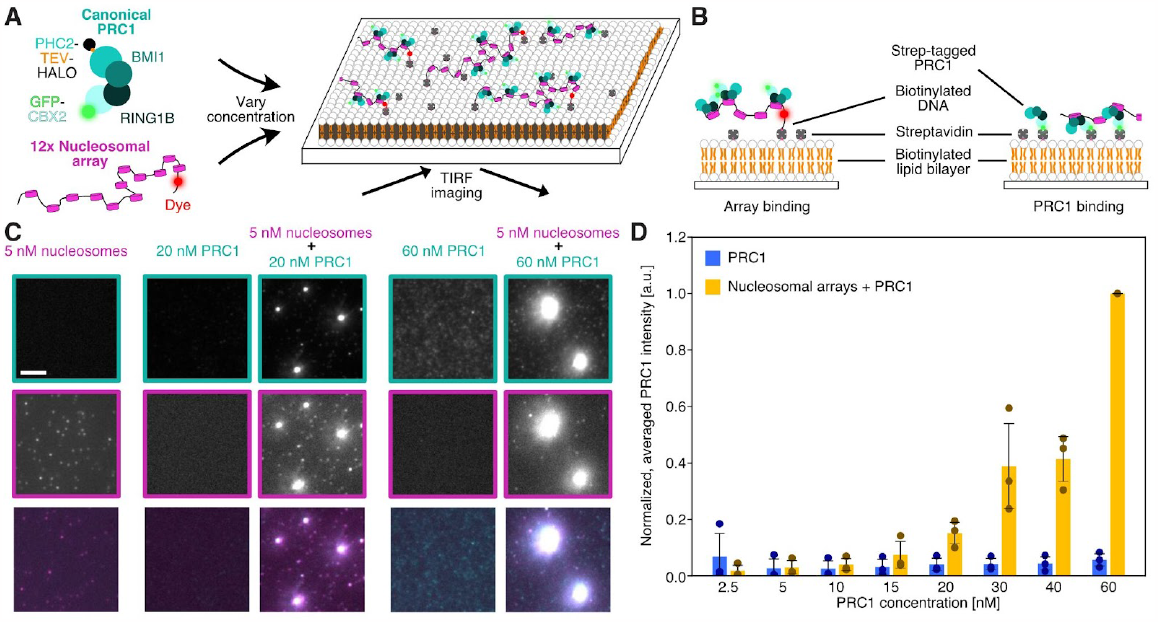
Nucleosomal arrays reduce critical concentration for PRC1 condensate formation. (**A**) Schematics of TIRF microscopy assay to study condensate formation of nucleosomal arrays and canonical PRC1 (for more details see **Figure S5A, B**). (**B**) Close-up view showing how nucleosomal arrays or PRC1 are attached to the biotinylated lipid bilayer. Left: 3’ and 5’ end biotinylated nucleosomal arrays are bound to biotinylated lipids via streptavidin and PRC1 is floating freely. Right: StrepII-tagged BMI of the full PRC1 complex is bound to biotinylated lipids via streptavidin (see **Materials and Methods** for details). (**C**) Representative TIRF microscopy images of nucleosomal arrays (magenta), PRC1 (cyan), and when both are mixed at various concentrations. The scale bar is 5 μm. (**D**) Averaged PRC1 intensities of all spots for PRC1 alone (blue) or for PRC1 mixed with nucleosomal arrays (orange). The nucleosome concentration was 5 nM. Dots represent the average PRC1 intensity from each of the three technical replicates. For each replicate 100 to >1000 spots were analyzed. Error bar is standard error of the mean.

We first asked if PRC1 alone, nucleosomal arrays alone, or a mixture of both will form condensates at low nanomolar concentrations. To test this, we imaged these different conditions after a 30-minute incubation at 30°C using the TIRF assay. We found that neither nucleosomal arrays alone with a 5 nM nucleosome concentration nor PRC1 at 20 or 60 nM concentrations formed condensates (**Figure 6C**). However, when we combined nucleosomal arrays and PRC1 at similar concentrations as above, we observed condensates (**Figure 6C**). These condensates showed higher fluorescence intensities with increasing PRC1 concentration. Measuring the fluorescence intensities of PRC1-containing spots at 60 nM PRC1 revealed ∼20-fold higher levels in the presence of nucleosomal arrays than for PRC1 alone (**Figure 6D**). We confirmed that this observation held true at the high nanomolar concentrations and imaged PRC1 alone, nucleosomal arrays alone, or a mixture of both with 480 nM PRC1 and 20 nM nucleosome using epifluorescence microscopy and only observed condensates when PRC1 and arrays were both present in the same chamber (**Figure S5B**). Taken together, we conclude that PRC1 and nucleosomal arrays act synergistically to form condensates, such that stable condensates form even at PRC1 concentrations of tens of nanomolar.

### Saturation of nucleosomal arrays with PRC1 is a prerequisite for condensate formation

We explored the concentration dependence for PRC1 condensate formation with nucleosomal arrays by systematically varying PRC1 concentration from 0 to 60 nM while keeping the nucleosome concentration constant at 5 nM and used the TIRF microscopy assay with biotinylated nucleosomal arrays as described above. We observed that the radius and brightness of condensates increased substantially with increasing PRC1 concentration for complexes containing either PHC2 WT or PHC2 L307R (**Figure 7A, Movie S5**). To quantitatively determine the PRC1-concentration dependence for condensate formation, we measured PRC1 fluorescence intensities of PRC1 bound to single nucleosomal arrays and in condensates. At PRC1 concentrations lower than the nucleosome concentration, the distribution of PRC1 intensities revealed a single population with increasing mean intensity as a function of PRC1 concentration (purple marker in **Figure 7B**). This indicates an increased binding of PRC1 to nucleosomal arrays with increasing PRC1 concentration. Interestingly, the intensity distribution broke into two populations (bimodal distribution) when the PRC1 concentration surpassed the nucleosome concentration (blue and orange marker in **Figure 7B**). The population with the lower PRC1 intensities (blue marker) remained at an almost constant mean intensity even when the PRC1 concentration was increased while the mean intensity of the population with the higher PRC1 intensities (orange marker) kept growing until it was saturated (**Figure 7B, C**). We also observed this bimodal distribution for nucleosomal arrays (**Figure S5C, D**). By converting the PRC1 and nucleosomal array intensities to the number of molecules based on photobleaching steps, we determined that there are roughly 10-20 PRC1 molecules per 12-mer nucleosomal array in the lower PRC1 intensity population while we measured around 10,000 PRC1 molecules in the population with higher PRC1 intensity (**Figure S5E, F**). Thus, a 12-fold increase in PRC1 concentration led to a ∼1,000-fold increase in the number of PRC1 molecules. Taken together, we conclude that at PRC1 concentrations lower than the nucleosome concentration no condensates form and that at PRC1 concentrations higher than the nucleosome concentration two populations emerge. We refer to the lower intensity population as “Saturated array” and to the higher intensity population as “Condensate” (**Figure 7B**). This bifurcation of the bimodal distribution is consistent with a switch-like transition for which nucleosomal arrays must first be saturated with PRC1 before condensates emerge and grow in size. This behavior is reminiscent of surface condensation or the prewetting phenomena which has recently been described for other protein-chromatin interactions^53^.

**Figure 7.**
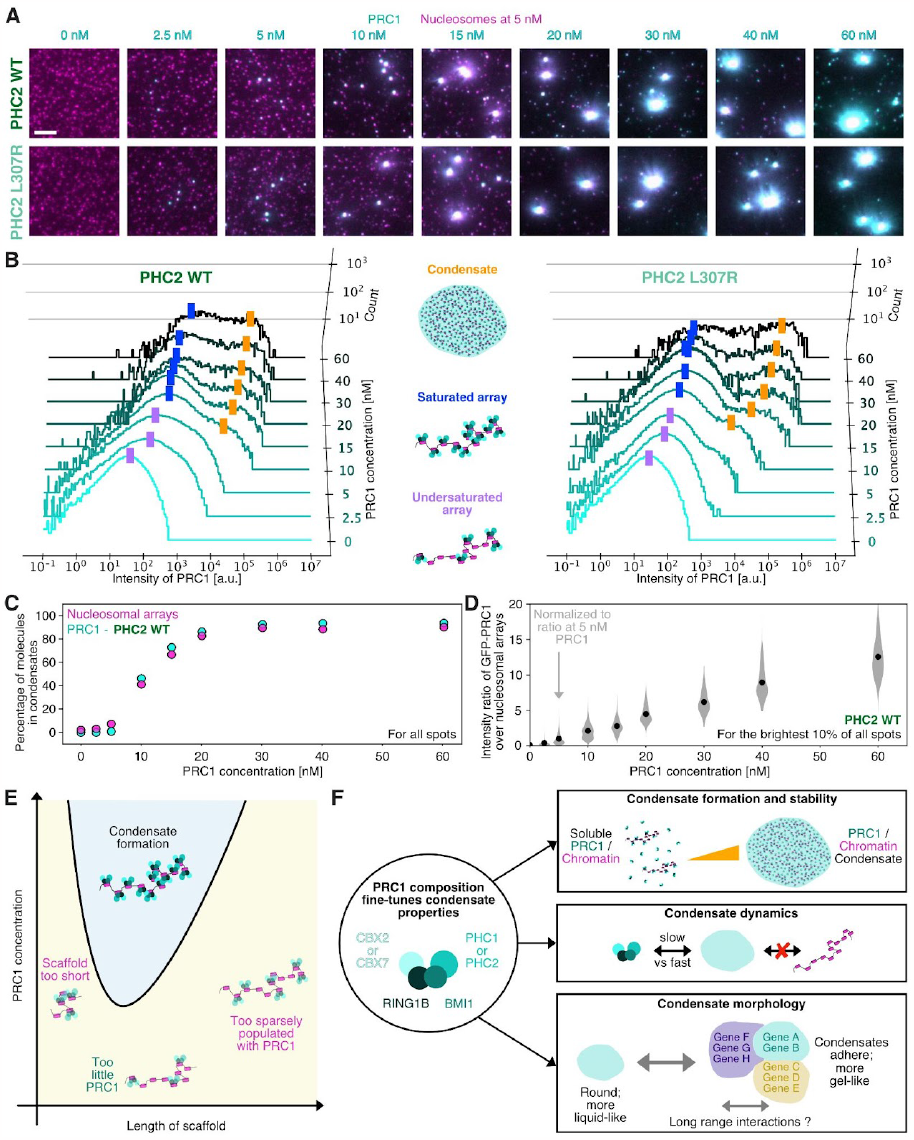
Stoichiometric excess of PRC1 to nucleosomes drives condensation. (**A**) Representative TIRF micrographs of PRC1 (cyan) and nucleosomal arrays (magenta) on lipid bilayers with either PHC2 WT or L307R. The scale bar is 5 μm. (**B**) Histograms of PRC1 intensities of spots as seen in **A** for PRC1 complexes with PHC2 WT (left) and PHC2 L307R (right). The purple rectangles indicate spots with undersaturated nucleosomal arrays, the blue rectangles correspond to single, saturated nucleosomal arrays, and the oranges rectangles show condensates of nucleosomal arrays and PRC1. (**C**) Percentage of molecules in condensates of PRC1 (cyan) and nucleosomal arrays (magenta) for PHC2 WT as shown in **B**. (**D**) Violin plot of intensity ratio of PRC1 over arrays normalized to the average ratio at 5 nM PRC1 for the brightest 10% of all spots as shown in **B**. (**B-D**) Data are representative of at least three technical repeats. (**E, F**) Model for how condensates of nucleosomal array / chromatin and PRC1 might form and how PRC1 composition determines condensate properties. Model is described in detail in the **Discussion**.

Based on the number of PRC1 molecules bound to saturated nucleosomal arrays and in condensates, we estimated the percentage of PRC1 molecules and nucleosomal arrays in condensates as a function of PRC1 concentration. For both, PRC1 and nucleosomal arrays, we found a sigmoidal distribution with a Hill coefficient of ∼5 suggesting a cooperative effect (**Figure 7C**). Furthermore, we measured that the ratio of PRC1 bound to nucleosomes is roughly one when they are present at equimolar concentrations and do not form condensates, but that the PRC1 concentration in condensates is up to 20-fold higher than the nucleosome concentration (**Figure 7D**). We conclude that condensates at the nanomolar regime will only emerge if PRC1 is in stoichiometric excess to nucleosomes. Together, the data suggests that condensate formation is driven by the cooperativity of PRC1-PRC1 interactions, but that these condensates are only stable because of PRC1 interactions with the nucleosomal array scaffold.

We validated our conclusions in multiple ways. First, we asked whether HALO-tag cleavage alters the outcome of PRC1 and nucleosomal array condensate formation. To this end, we expressed and purified a PRC1 complex containing mGFP-CBX2, RING1B, BMI1, and PHC2 L307R without the C-terminal HALO-tag (**Figure S5G**). With this PRC1 construct, we performed the same TIRF microscopy assay as described above and observed the same bimodal distribution as we saw for PHC2 WT and L307R with the cleavable HALO-tag (**Figure S5H, Movie S5**). This suggests that HALO-tag cleavage does not alter condensate formation. Next, we validated that our observations were not caused by the lipid bilayer by using an alternative biotin-PEG surface treatment and found very similar results indicating that the condensate formation is not a surface artifact (**Figure S5I-K)**. Third, instead of mixing PRC1 and nucleosomal arrays in a test tube for 30 min before adding them to the bilayer, we also imaged condensate formation in real time right after mixing nucleosomal arrays and PRC1. In agreement with our previous results, this real time imaging showed that condensates only appeared if the PRC1 concentration was higher than the nucleosome concentration and that the intensity distributions of PRC1 followed the previously described bifurcation, which emerged within minutes after starting the experiment (**Figure S6A-E**). Fourth, we validated that the stoichiometric excess of PRC1 to nucleosomes is important for condensate formation by also altering the nucleosome concentration. We found that with increasing nucleosome concentration a higher PRC1 concentration is required for condensation (**Figure S6F, G**). Furthermore, we found that adding more nucleosomal arrays to pre-formed PRC1-nucleosome condensates can shift the equilibrium and reduce the intensity of condensates (**Figure S6H-J**). Both observations support our conclusion that the ratio of PRC1 to nucleosome concentration is critical for condensate formation and maintenance. Taken together, we conclude that PRC1 and nucleosomal arrays act synergistically to form condensates and that condensation is driven by excess PRC1 to nucleosomal arrays so that a prewetted chromatin state is created which in turn can grow into condensates by more PRC1 binding.

### Length of nucleosomal arrays is critical for condensate formation

Since the saturation of nucleosomal arrays with PRC1 proved to be important for condensate formation, we asked whether specific nucleosomal array properties such as array length, nucleosome density, or histone modifications play a role in condensation. We first tested the scaffold model by changing the length of nucleosomal arrays. In addition to the 12mer 5S G5E4 array that we used so far, we added 12mer, 4mer, and 1mer arrays based on the Widom 601 sequence^54^ (**Figure S7A**). For the tetra- and mononucleosomes we kept either the DNA concentration or the nucleosome concentration constant with respect to the 12mer nucleosomal array. Analyzing the intensities of our images, we measured a correlation between nucleosomal array length and condensation (**Figures S7B, C**). For instance, at 20 nM PRC1 concentration and constant nucleosome concentration of 5 nM we determined that 8, 20, or 40 percent of PRC1 molecules are in condensates for 1mer, 4mer, or 12mer arrays, respectively. We next varied the nucleosome density on arrays by varying linker length and measured only small differences (**Figure S7D-F**). Furthermore, we observed that DNA alone in the absence of nucleosomes is ∼7-fold weaker in forming condensates than nucleosomal arrays at 20 nM PRC1 suggesting that PRC1 binding to nucleosomes is critical to promote condensation (**Figure S7G, H**). Lastly and in agreement with other studies^18,50,55^, we did not observe any significant differences for arrays with unmodified histones compared to arrays with histone modifications such as H3K27me3 (**Figure S7I, J**). We conclude that nucleosomal arrays act as a scaffold and that scaffold length and presence of nucleosomes are important determinants for lowering the critical concentration for PRC1 condensation.

## Discussion

Polycomb repressive complexes are key to maintaining repression of hundreds of genes and are essential for proper development^7–12^. The ability of canonical PRC1 to form condensates *in vitro* and in cells has been suggested to contribute to maintenance of repression. In this study, we discovered that chromatin and PRC1 act synergistically to form condensates leading to a significant decrease of the critical concentration required for PRC1 condensate formation. Using single-molecule and live cell imaging, we found that the exact combination of CBX and PHC subunits in the PRC1 complex defines initiation, morphology, stability, and dynamics of condensates. Mutations in CBX2 (23KRA) and PHC2 (L307R) that are known to impact mouse development substantively alter condensate formation^14,44^. We propose that the spectrum of PRC1 condensate properties is a mechanism through which PRC1 determines the extent of gene repression to impact developmental progression.

Specifically, we propose the following model for the formation and regulation of chromatin and PRC1 condensates (**Figure 7E, F**). First, the chromatin scaffold must be of a sufficient length and nucleosomes must be densely bound by PRC1. Second, PRC1 must be in stoichiometric excess to nucleosomes for condensates to form. Third, the combination of the CBX and the PHC subunit of PRC1 must have the right properties to initiate condensate formation. For instance, PRC1 with CBX2 forms larger condensates compared to PRC1 with CBX7 which will only form condensates if PHC1 but not PHC2 is present. Our model further suggests that once these requirements are met and large enough condensates have formed, that PHC is critical for generating and stabilizing distinct, long-lived domains that can adhere to each other but do not coalesce. The adherence of different domains might in turn be required for the formation of long-range interactions and Polycomb bodies. Below, we will discuss the details and biological implications of our model in the context of previous findings.

### Chromatin scaffold model explains regulation of PRC1 condensate formation

In this study we observed a dual-component scaffold mechanism for the condensate formation of chromatin and PRC1 (**Figure 7E**). We show that nucleosomal arrays enable condensation of PRC1 at concentrations far below the saturation concentration for condensate formation of PRC1 alone (**Figure 6**). The requirement of a scaffold to drive condensate formation has also been observed for other biological processes^28,55–62^. For instance, Shrinivas et al.^28^ found that DNA motif valency and density drive the formation of transcription factor condensates and that the ratio of client to scaffold is important to initiate condensation. This example is in good agreement with our observations for PRC1 as we also observed a dependency of condensate formation on scaffold length, scaffold valency, and ratio of scaffold to client (**Figures 7, S7**). Together with other studies^28,55–61^, this suggests that similar biophysical principles in the nucleus might be used to organize activated and repressed genes. In these examples and our observations, chromatin or DNA serve as scaffold and its organization by its various clients depends on meeting key thresholds. Such a tightly regulated scaffold mechanism might explain how condensate formation is restricted to specific genomic loci as condensates will only emerge if the key parameters such as scaffold length and/or scaffold to client ratio have been met.

The single-molecule imaging protocol we developed revealed a sigmoidal behavior for condensate formation. Specifically, we observed that only nucleosomal arrays saturated with PRC1 underwent a switch-like transition to form condensates (**Figures 7, S5, S6**). This observation agrees well with another study in which a DNA scaffold must be fully coated with the pioneer transcription factor Klf4 before condensates form^53^. Specifically, Morin et al. showed that Klf4 condensates on DNA as a type of surface condensation (prewetting transition). For PRC1 and chromatin, this surface condensation model suggests that PRC1 molecules can independently read out histone information and bind nucleosomes while collectively integrating histone information into the condensed state. In this model, the chromatin scaffold acts as a signal amplifier and provides a more robust mechanism to regulate gene transcription. Taken together, condensate formation via a scaffold-client mechanism is emerging as a general principle to organize chromatin structure and to restrict it to well-defined loci in the nucleus.

The FRAP and two-color mixing experiments (**Figure 2**), in agreement with live cell imaging data^63,64^ suggest that there are two different classes of PRC1 molecules present in condensates: one PRC1 class that is tightly bound to chromatin and one that is more diffusive. Interestingly, these live cell imaging experiments from Huseyin et al.^63^ estimated the PRC1 concentration in Polycomb bodies to be 130 nM while condensate formation of PRC1 *in vitro* has previously been observed at the low micromolar and high nanomolar concentrations^18,19^. This raises questions whether there is sufficiently high PRC1 concentration to form condensates in the nucleus. However, we found that PRC1 together with nucleosomal arrays can form condensates at low nanomolar concentrations (∼10 nM) without crowding reagents (**Figures 6, 7**), indicating that condensation of PRC1 in the nucleus is possible with the estimated nuclear concentration of PRC1. Thus, with the single-molecule TIRF imaging approach we have reconciled some previous discrepancies between live cell and *in vitro* data showing that PRC1 and chromatin condensate formation is indeed possible under conditions present in the nucleus.

### Roles of CBX and PHC subunits in condensation and gene repression

The single-molecule and live cell imaging data demonstrated that the complex composition of the PRC1 complex plays a critical role for the formation of condensates. In particular the specific combination of the two families of subunits that have previously been shown to form condensates, CBX and PHC^18,19,38^, determine whether condensates form, and influence the morphology, stability, and dynamics of condensates (**Figure 7F**).

While CBX2 forms condensates with either PHC1 or PHC2, we found that CBX7 only forms condensates when PHC1 is present in the same PRC1 complex (**Figure 4**). This offers an explanation for why CBX7 is found in condensates in cells despite it lacking any intrinsic ability to form condensates. The nature of these condensates might contribute to the greater plasticity of the pluripotent and multipotent cells that express CBX7 and PHC1^13,41^. As described above and seen in previous work^19^, CBX2 condensation is driven by the CaPS region. Mutations in the highly positively charged region of the CBX2 CaPS region diminish condensate formation. We found in this study that PHC1 can drive condensate formation independent of the properties of the CBX subunit, in contrast to PHC2 which only forms condensates when a CBX protein capable of forming condensates, such as CBX2, is present in the same PRC1 complex (**Figures 3, 4, 5**). Hence, PHC1 appears to have a stronger propensity to initiate condensate formation than PHC2. We found that the N-terminal part of PHC1 is important for this process as the C-terminal part of PHC1 alone cannot form condensates with CBX7. Interestingly, previous studies have shown that the N-terminus of the *Drosophila* PHC protein (Ph) can be modified by the glycosyltransferase Ogt by adding O-linkedN-Acetyl-glucosamine (O-GlcNAc) and suggested that this modification is important for the ordered assembly of PRC1 molecules^52,65,66^. Together with data showing that CBX2 phosphorylation by casein kinase II (CK2) is important for condensate formation^19,67^, this suggests that in addition to the complex composition posttranslational modifications can further fine-tune and regulate PRC1 condensate properties.

Our work revealed that the specific subunit composition of PRC1 also influences the morphology of condensates and that this is mostly dependent on the properties of the PHC subunit.

For instance, PRC1 condensates with PHC2 wild type are elongated while condensates with polymerization deficient PHC2 are perfectly round suggesting that the polymerization activity of the SAM domain of PHC2 is responsible for this morphology change (**Figure 1**). Interestingly, super-resolution imaging data of a study that investigated the long-range interactions of *Drosophila* Ph also showed these morphology differences for PRC1 with active and inactive polymerization activity^15^. While it is challenging to understand the molecular mechanism behind these morphology changes in cells, by using the reconstituted system we were able to investigate the nature of the modularity in PRC1 interactions. We used multi-color colocalization experiments to show that the morphology changes arise from many, small condensates of PRC1 and chromatin that adhere but not coalesce (**Figure 2**). One possible explanation for these changes is that condensates with PHC2 polymerization undergo a transition from more liquid-like behavior to a more gel-like state as shown by FRAP analysis (**Figure 2**). Together with data from previous studies^13–15,68^, we propose that the phase transition that leads to the adherence of condensates without coalescence might play an important role in driving long-range chromatin interactions and in organizing chromatin into different domains. In this way many smaller condensates can be brought together into larger domains (e.g. Polycomb bodies) without fusion (**Figure 7F**). This might be biologically advantageous as it would allow for the maintenance of many genes together in one locus, while simultaneously keeping them separate. Interestingly, adherence without fusion is not unique to PRC1 and chromatin, but has also been observed for an artificial chromatin-based system^25^ suggesting that this might be a more broadly employed mechanism to compartmentalize chromatin in the nucleus.

The changes in PRC1 condensate properties that lead to adherence without coalescence also provide an explanation for the changes that we see in PRC1 condensate stability and dynamics. For instance, we found that PRC1 with WT PHC2 or PHC1 are more robust to salt changes *in vitro* indicating that these condensates are more stable than complexes with polymerization deficient PHC2 (**Figure 1**). Indeed, live cell imaging revealed that the number of visible CBX2 condensates is dramatically reduced if a polymerization deficient version of PHC2 is co-expressed suggesting that CBX condensation alone is not sufficient to maintain condensates in cells (**Figure 5**). This observation is in good agreement with previous work which showed that RING1B puncta disappear if a PHC polymerization deficient construct is expressed in cells^14^. The increased stability by PHC polymerization is likely because more PRC1 molecules are retained in condensates as shown by the FRAP data (**Figure 2**). Together with structural data of PHC polymerization^46^, we suggest the polymerization of the SAM domain provides a second scaffold in addition to the chromatin scaffold, which increases the retention rate of PRC1 and might make PRC1 condensates more robust and enhance repression. Taken together, the exact combination of CBX and PHC subunits in PRC1 complexes is important for condensation and neither subunit can work alone to create and maintain condensates. Since both subunits have many paralogs and in particular CBX paralogs have been shown to have a wide range of compaction and condensation capabilities, that can change gene expression patterns^18,19,43,44,50,69^, it is possible that the various compositions of PRC1 might offer important flexibility of regulatory function during different stages of development.

## Acknowledgments

We are grateful to Jongmin J. Kim, Wojciech Siwek, MacKenzie Mauger, and all members of the Kingston and Subramanian laboratories (all Massachusetts General Hospital Research Institute and Harvard Medical School, Boston) for critical discussions of this work. We thank Richard Bouley from the Massachusetts General Hospital Program in Membrane Biology (PMB) Microscopy Core for support with the FRAP and live cell data collection. We thank Luke H. Chao and Melody Nguyen for advice on the lipid bilayer system. S.N. is the Dennis and Marsha Dammerman Fellow of the Damon Runyon Cancer Research Foundation (DRG-2418-21). The authors gratefully acknowledge funding from NIH grant R35GM131743 to R.E.K..

## Author Contributions

S.N., R.S., and R.E.K. designed the research; S.N. and S.M. cloned constructs and prepared baculovirus. S.M. designed and validated the activatable PHC system. S.N. expressed and prepared samples; T.A.O. generated dox-inducible 3T3 cell lines; S.N. collected TIRF, epi, and confocal microscopy data; S.N. developed data collection and analysis code; S.N. analyzed the data; S.N., R.S., and R.E.K. wrote the manuscript. All authors read and commented on the paper.

## Materials and Methods

### Protein expression, purification, and labeling

All PRC1 complexes were expressed in Sf9 cells using the Bac-to-Bac system (Thermo Fisher Scientific, 10359016) as previously described^19^. We always expressed all four subunits of canonical PRC1 (RING1B, BMI1, CBX, and PHC) together. For RING1B, and BMI1, we cloned the full length, untagged mouse cDNA into a pBig1a vector^70^ (Addgene, Plasmid #80611). These untagged RING1B and BMI1 constructs were used for all experiments presented in this manuscript with the exception for the data presented in **Figure 6C, D**. Here, we used StrepTagII-GSA-BMI1 to bind the full canonical PRC1 complex alone (or with non-biotinylated nucleosomal arrays) to biotinylated lipid bilayers via streptavidin (**Figure 6**). We chose the StrepTagII instead of direct biotinylation of BMI1 via for instance a BirA-tag to avoid overcrowding of the bilayer surface with single PRC1 molecules. Since the StrepTagII has a weaker affinity for Streptavidin, we could enrich the binding of samples that had more than one or two PRC1 molecules and thus reduce the background of single PRC1 molecules.

For the CBX subunits (mouse), we cloned the following constructs into a pFastBac1 vector (Thermo Fisher Scientific, 10359016): Flag-GSAAAGS-mGFP-GSAAAGS-CBX2, Flag-HALO-GSAAAGS-CBX2, Flag-GSAAAGS-mGFP-GSAAAGS-CBX2-13KRA, Flag-GSAAAGS-mGFP-GSAAAGS-CBX2-23KRA, Flag-GSAAAGS-mGFP-GSAAAGS-CBX7, and Flag-GSAAAGS-mGFP-GSAAAGS-CBX7-CaPS. For PHC subunits (mouse), we cloned the following constructs into pFastBac1 vector (Thermo Fisher Scientific, 10359016): PHC2-L307R, PHC2-TEV-HALO, PHC2-L307R-TEV-HALO, StrepTagII-GSAAAGS-TEV-PHC1-TEV-HALO, and StrepTagII-GSAAAGS-TEV-PHC1-L994R-TEV-HALO.

We infected Sf9 cells (Expression Systems, 94-001F) with a multiplicity of infection of 5 of baculovirus at 1×10^6^ cells per ml and incubated at 27°C for 70 hours in Protein-free Insect Cell Culture Medium (Expression Systems, 96-001-01) with penicillin/streptomycin. Cells were harvested by centrifugation at 5,000 RCF for 20 min. The cell pellet of a 1-liter expression was resuspended in 20 ml of PBS, pH 7.4. Afterwards, cells were spun again at 1,000 RCF for 5 min and the supernatant was discarded. We next measured the packed cell volume (PCV) and resuspended the pellet in a hypotonic buffer (10 mM HEPES at pH 7.9, 10 mM KCl, 1.5 mM MgCl_2_, 0.1 mM DTT, 0.2 mM PMSF) to 3x PCV. The cells were incubated on ice for 10 min and lysed with a Dounce homogenizer. This was followed by a centrifugation step at 4,000 RCF for 15 min. Subsequently, we measured the packed nuclear volume (PNV) of the pellet and resuspended the nuclei in a volume of low salt buffer (20 mM HEPES at pH 7.9, 20 mM KCl, 1.5 mM MgCl_2_, 0.2 mM EDTA at pH 8.0, 25% glycerol, 0.1 mM DTT, 0.2 mM PMSF) equal to 1/2 PNV. Afterwards, we added a volume of high salt buffer (20 mM HEPES at pH 7.9, 1200 mM KCl, 1.5 mM MgCl_2_, 0.2 Mm EDTA at pH 8.0, 25% glycerol, 0.1 mM DTT, 0.2 mM PMSF) equal to 1/2 PNV and incubated the solution with gentle mixing at 4°C for 30 min. The nuclei were pelleted by centrifugation at 20,000 RPM for 30 minutes (25,000 RCF). Finally, we flash-froze the supernatant (nuclear extract) and stored it at −80°C.

The nuclear extract was subsequently incubated with MonoRab™ Anti-DYKDDDDK affinity resin (Genscript, L00766) for 2 hours and then washed with BC300 buffer (20 mM HEPES at pH 7.9, 300 mM KCl, 1 mM EDTA, 1 mM MgCl2, 10% glycerol, 0.05% IGEPAL CA-630 (Sigma, I8896-100ML), 1 mM DTT, 0.1 mM PMSF, cOmplete EDTA-free protease inhibitor [Roche]). Afterwards the beads were washed with BC600 buffer (20 mM HEPES at pH 7.9, 600 mM KCl, 1 mM EDTA, 1 mM MgCl2, 10% glycerol, 0.05% IGEPAL CA-630, 1 mM DTT, 0.1 mM PMSF, cOmplete EDTA-free protease inhibitor [Roche]) and again with BC300 buffer without IGEPAL CA-630. PRC1 complexes were eluted from the resin using BC300 buffer (without IGEPAL CA-630) containing 0.8 mg/mL Flag peptide.

For PRC1 complexes containing PHC1, we added an additional purification step and incubated the Flag-eluted samples with Strep-TactinXT 4Flow beads (IBA lifesciences, 2-5030-010) for 2 hours. Then the resin was washed with a BC300 buffer without IGEPAL CA-630. Protein was eluted from the resin using BC300 buffer (without IGEPAL CA-630) supplemented with 50 mM Biotin.

Subsequently all PRC1 complexes were concentrated using Amicon Ultra-4 centrifugal filter units with a MW cutoff of 100 kDa (MilliporeSigma, UFC810024). If applicable, we labeled HALO-tagged^49^ proteins with HALO-Alexa488 or HALO-TMR dyes (Promega, G1001, G8251) by incubating the concentrated protein with dye for 1 hour on ice. Afterwards, PRC1 complexes were further purified by size exclusion chromatography on a Superose 6 Increase 10/300 GL column (Cytiva, 29-0915-96). Fractions containing full PRC1 complexes were concentrated using Amicon Ultra-4 centrifugal filter units with a MW cutoff of 100 kDa (MilliporeSigma, UFC810024) and protein concentration was determined by Bradford assay. Finally, we confirmed the purity of complexes by 4-20% SDS-PAGE (Biorad, #4561096) and Coomassie staining. Protein aliquots were flash-frozen and stored at −80°C.

### Preparation of nucleosomal arrays

Recombinant histones were purified based on a protocol from Dyer et al^71^. The recombinant Xenopus histones H2A, H2B, H3, and H4 in a pET28 vector were expressed in BL21 DE3 cells and grown at 37°C in 2YT with 50 mg/ml kanamycin. After induction with 1mM IPTG, cells were grown for 3 more hours. Then the cells were harvested by spinning at 5,000 RCF for 20 minutes. Subsequently, the cell pellets were resuspended and sonicated in a wash buffer (50 mM Tris at pH 7.5, 100 mM NaCl, 1 mM PMSF, and 1 mM 2-mercaptoethanol). The inclusion bodies were pelleted at 48,000 RCF for 15 minutes. Then pellet was resuspended in wash buffer containing 1% Triton X-100 and pelleted again at 48,000 RCF for 15 minutes. This wash step was repeated twice, and then the pellet was resuspended in SAUDE 200 (7 M urea, 20 mM sodium acetate at pH 5.2, 200 mM NaCl, 5 mM 2-mercaptoethanol, 1 mM EDTA). Afterwards, the sample was centrifuged at 48,000 RCF for 15 minutes and the supernatant was filtered through a 0.22-micron syringe filter (GenClone, 25-243). The sample was loaded onto a 5 mL HiTrap SP HP column (Cytiva, 29-0513-24) equilibrated in SAUDE 200, washed with 10% SAUDE 1000 (7 M urea, 20 mM sodium acetate at pH 5.2, 1000 mM NaCl, 5 mM 2-mercaptoethanol, 1 mM EDTA), and eluted with a linear gradient from 10% to 50% SAUDE 1000. The histone fractions were dialyzed overnight into 5 mM 2-mercaptoethanol and lyophilized.

The plasmids for expressing acetylated histones (pCDF-pylT H3 and pBK-AcKRS) were a kind gift from Jason Chin. Plasmid pCDF-pylT H3K27ac was cloned by mutating the lysine 27 codon in plasmid pCDF-pylT H3 to an amber stop codon. Plasmids pCDF-pylT H3K27ac and pBK-AcKRS were transformed into BL21 (DE3) cells and grown overnight in 2YT media supplemented with 30 μg/ml streptomycin and 50 μg/ml kanamycin. At an OD600 of 0.7–0.8, the culture was supplemented with 20 mM nicotinamide (NAM) and 10 mM acetyl-lysine (AcK). Protein expression, purification, dialysis, and TEV cleavage were performed as described by Neumann et al.^72^ with the following changes. Extracted protein was bound to Ni^2+^ resin (Ni Sepharose 6 Fast Flow, GE Healthcare) in 6M guanidine HCl, 20 mM Tris (pH 7.5), 200 mM NaCl. The column was washed with 6M urea, 20 mM Tris (pH 7.5), 200 mM NaCl, 25 mM imidazole and bound protein was eluted with 6M urea, 20 mM Tris (pH 7.5), 200 mM NaCl, 1M imidazole. Following dialysis and TEV cleavage, H3K27ac was lyophilized and stored at −80°C. The methylated Xenopus H3 histones (H3K4me3, H3K27me3) were purchased from The Histone Source at Colorado State University (XH3_K4me3, XH3_K27me3).

The recombinant histone octamer was assembled using the histones prepared as described above, with 4 mg of each histone dissolved in 2 mL unfolding buffer (6 M guanidinium chloride, 20 mM TRIS at pH 7.5, 5 mM DTT). Afterwards, the histones were combined in equimolar amounts and dialyzed in Slide-A-Lyzer Dialysis Cassettes, 7 kDa (Thermo Fisher Scientific, 66370) overnight at 4°C against three changes of 1 L refolding buffer (2 M NaCl, 10 mM TRIS at pH 7.5, 1 mM EDTA, 5 mM 2-mercaptoethanol). Precipitated protein was removed by centrifugation at 48,000 RCF for 15 minutes. The sample was then loaded onto a HiLoad 26_600 Superdex 200 gel filtration column (Cytiva, 28-9893-36) equilibrated with refolding buffer. Octamer fractions were combined and stored at 4°C.

The DNA for nucleosomal arrays were cloned into pUC57 vectors. We modified the G5E4 plasmid^47,48^, by replacing the ClaI and KpnI restriction sites with BsmBI. The BsmBI sites allowed for the usage of nonpalindromic sequences so that other dsDNA can be ligated to the array strands without getting self-ligation of the array strands. DNA containing the Widom 601 sequence^54^ to make dodeca-, tetra-, and mononucleosomes had a 20 bp linker between the 601 repeats. Here, we replaced the EcoRI and HindIII sites with BsmBI. The plasmids with various linker length were a gift from Michael Rosen^25^ (Addgene: pWM_12×601_15bpLinker #157786, pWM_12×601_20bpLinker #157787, pWM_12×601_25bpLinker #157788, pWM_12×601_30bpLinker #157789, pWM_12×601_35bpLinker #157790) and were modified by adding BsmBI just outside of the original EcoRV cutting sites. These plasmids were transformed into NEB 5-alpha Competent E. coli (New England Biolabs, C2987H) cells and grown with 100 μg/mL carbenicillin in TB (terrific broth). After harvesting the cells by centrifugation at 5,000 RCF for 15 minutes, the DNA was purified following manufacturer’s instructions for the Qiagen Plasmid Maxi Kit (Qiagen, 12162).

To separate the nucleosomal array DNA from the vector backbone we performed the following steps: first, 8.6 ml of DNA at 10 mg/ml were mixed with 1.0 ml of 10x CutSmart buffer, 0.2 ml of BsmBI_v2 (NEB, R0739L) and incubated at 55°C for 48 hours. Afterwards, we added 0.2 ml of DdeI (NEB, R0175L) for the 2×601, 12×601, and G5E4 DNA or 0.2 ml of AvaII (NEB, R0153L) for the 12×601 with various linker lengths and incubated at 37°C for 24 hours (except for the Tetra-nucleosome 601 DNA). We then ran 2 μl of the sample on a 1% agarose gel to check if the reaction was completed. If necessary, additional enzyme(s) was added and incubated for another 12 hours at the respective temperature. Next, we added 5 M NaCl to the DNA for a final concentration of 0.5 M NaCl. Afterwards, a PEG precipitation was performed (except for the Tetra-nucleosome 601 DNA). To this end, PEG-6000 (Sigma, 8074911000) with 0.5 M NaCl was added to ∼5% final (depending on sample) and incubated at 4°C overnight. The next day, the sample was spun at 29,600 RCF for 20 minutes. The pellet was resuspend in 7 ml of TE buffer (10 mM TRIS pH at 8.0, 1 mM EDTA at pH 8.0), and allowed to rehydrate at 4°C for 4 hours. For the Tetra-nucleosome 601 DNA we used electroelution to separate the backbone from the DNA of interest. To this end, all DNA was run on a 1% agarose gel, the band of interest was excised and placed in a dialysis bag (Spectrum Labs, 133192) with TBE. Then, the dialysis bag was submerged in a gel chamber with TBE Buffer and DNA eluted for 1 hour at 80V on ice. Afterward, the solution was pipetted out of the dialysis bag. Subsequently, the DNA was purified by phenol:chloroform:isoamyl alcohol (25:24:1) extraction and ethanol precipitation and the DNA was resuspended in TE buffer.

To label the DNA with fluorophores or biotin, we ordered oligonucleotides from Integrated DNA Technologies (IDT): OligoA-BsmBI_Side1_Biotin (Biotin/ TTT TTT TTG GTG TAG GAG GTA GAT GAG); OligoA-BsmBI_Side1_UN (TTT TTT TTG GTG TAG GAG GTA GAT GAG); OligoB-BsmBI_Side1_ATTO647N or Cy3N (Phos ATCG CTC ATC TAC CTC CTA CAC CAA /ATTO647NN/); OligoA-BsmBI_Side2_Biotin (Phos ACTG GGT AGA GTG GTA AGT AGT GAA TTT TTT /Biotin/); OligoB-BsmBI_Side2UN (/Cy3N/ TTC ACT ACT TAC CAC TCT ACC). Oligos for side 1 and 2 were dimerized separately by mixing 9 μl of OligoA at 100 μM with 9 μl of OligoB at 100 μM and 2 μl of 10x DNA Ligase Buffer (NEB, M0202S) and heating to 95 °C for 5 minutes, followed by cooling down to 20°C at a rate of 1°C per minute. This created dsDNA fragments with complementary BsmBI sides to the digested nucleosomal array strands. These dsDNA fragments were then ligated to the nucleosomal array strands with T4 DNA ligase (NEB, M0202S) by incubating at 16°C overnight (dsDNA was used in 8-fold excess compared to array strands). The excess dsDNA was removed by 3 rounds of PCR clean up with QIAquick PCR Purification Kit (Qiagen, 28106).

Nucleosomal arrays were assembled as previously described^73^. Briefly, 5 μg of DNA and 5 μg of histone octamer were added to high-salt buffer (10 mM TRIS at pH 7.5, 2 M KCl, 1 mM EDTA, 1 mM DTT) in 7 kDa Slide-A-Lyzer MINI Dialysis Devices (ThermoFisher Scientific, 69562). Samples were placed in 400 ml high-salt buffer, and a ‘‘rabbit’’ pump was set up to pump 1600 ml low-salt buffer (10 mM TRIS at pH 7.5, 100 mM KCl, 1 mM EDTA, 1 mM DTT) into the dialysis beaker while removing buffer from the beaker at the same rate. Samples were dialyzed at 4°C overnight, then dialyzed against a storage buffer (10 mM TRIS at pH 7.5, 10 mM KCl, 1 mM EDTA, 1 mM DTT) for 4 hours. Dialyzed nucleosomal arrays were centrifuged at 15,000 RCF for 10 minutes at 4°C to remove aggregates. Arrays were digested by BsrFI-v2 (G5E4 array) or by AccI (601 sequences) (NEB, R0682S, R0161S respectively) for 2 hours at 30°C and run on a 1% agarose gel to assess the extent of nucleosome occupancy on the positioning sequences and finally stored at 4°C (**Figure S1D**).

### Preparation of lipid bilayers

The lipid bilayers were prepared based on previously described protocols^74–76^. Briefly, we cleaned glass vials (Thermo Fisher Scientifc, 14-955-331) and Hamilton glass syringe (Avanti, 610000-1EA) with Milli-Q water, 70% ethanol, Chloroform, 70% ethanol, Milli-Q water and let them air dry. We then dissolved 300 mg of 18:1 (Δ9-Cis) PC (DOPC) (Avanti, 850375P-500mg), 25 mg 18:1 PEG2000 PE (Avanti, 880130P-25mg), and 2.5 mg 18:1 Biotinyl Cap PE (Avanti, 870273P-25mg) in 25 ml of chloroform. We then prepared 2 ml aliquots, dried them with nitrogen and stored them at −20°C (sealed with High-Density Thread Sealant Tape). When ready to use, dried lipids were resuspended in 2 ml chloroform and stored at −20°C for several weeks.

To prepare a lipid mixture for use, we transferred ∼160 μl of lipids from the resuspended stock into a glass vial using a cleaned (Milli-Q water, followed by 70% ethanol and Chloroform) Hamilton glass syringe (Avanti, 610000-1EA), dried it with nitrogen and stored it in a desiccator overnight. The next day, 1 ml of PBS at pH 7.4 was added to dissolve the dried lipids, resulting in a 1 mg/ml mixture. The glass vial was then left at room temperature for at least 10 minutes to ensure complete suspension of the lipids. The mixture was then sonicated for 5 minutes in a water bath sonicator (Branson 3800). Meanwhile, the lipid extruder (Avanti, 610000-1EA) was assembled by washing the syringes with Milli-Q water, then 70% ethanol, and again with Milli-Q water. The 0.1 μm filter paper (Avanti, 610005-1EA) and support filters (Avanti, 610014-1EA) were wetted with Milli-Q water. After sonication, 5 freeze-thaws were performed by transferring the vial with the lipids from liquid nitrogen to ∼40°C water five times. Following this, the lipids were added to the extruder and pushed 21 times. The final lipid mixture was stored at 4°C.

### Flow-cell preparation

The flow-cells were assembled as previously described^77^. In brief, we used double-sided adhesive sheets (Soles2dance, 9474-08×12 − 3M 9474LE 300LSE) and cut out custom three-cell flow chambers using a paper cutter (Silhouette, Portrait 3). We then used the double-sided tape and assembled three-cell flow chambers together with glass slides (Thermo Fisher Scientific, 12-550-123) and 170 μm thick coverslips (Zeiss, 474030-9000-000). Prior to assembly of flow-cells, we cleaned the coverslips in a 5% v/v solution of Hellmanex III (Sigma, Z805939-1EA) at 50°C overnight and washed the coverslips extensively with Milli-Q water afterwards.

### Preparation of flow-cells with lipid bilayer system

The flow-cells for TIRF imaging were prepared in slightly different ways depending on the experimental design. For end-point assays, we typically mixed nucleosomal arrays (5 nM nucleosome concentration final unless otherwise noted) and various concentrations of PRC1 in TIRF Buffer (13 mM HEPES at pH 7.9, 100 mM KCl, 1.4 mM MgCl2, 0.25 mM EDTA, 0.9 mM ATP, 1% glycerol, 0.04 μg/μl BSA, 0.4 μg/μl kappa casein). We used (unless otherwise noted) G5E4 arrays labeled with one ATTO-647N dye and two biotins as described above. We incubated the 30 μl mixture at 30°C for 30 minutes in a thermocycler. In the meantime, we prepared the flow-cells with lipid bilayers. To this end, we added 13 μl of lipid bilayers to the flow-cells, incubated for 2 minutes and washed with 3x 13 μl of TIRF Buffer. We then added 10 μl of Streptavidin (Vector, SA-5000) and incubated for 2 minutes. This was followed by two washes with 13 μl of TIRF Buffer. Afterwards, we added 13 μl of the array/PRC1 sample and incubated for 5 minutes before imaging.

For live imaging of condensate formation, we prepared a “seed” mixture by mixing nucleosomal arrays (2.5 nM nucleosome concentration) and 5 nM of PRC1 in TIRF Buffer. This was incubated at 30°C for 30 minutes. In the meantime, we prepared the flow-cell with a lipid bilayer as described above. Once the “seed” mixture was added to the flow-cell and incubated for 5 minutes, we added a mixture of nucleosomal arrays (2.5 nM nucleosome concentration) and various concentrations of PRC1 in the TIRF Buffer and immediately started imaging (∼1 minute delay). For this additional mixture, PRC1 was added right before the solution was added to the flow-cell, to prevent condensate formation before imaging started.

For mixing arrays and PRC1 in different colors, we mixed nucleosomal arrays and PRC1 as described above making sure that we kept arrays or PRC1 in different colors separate. For differently colored PRC1, we mGFP-tagged the CBX subunit of the full canonical PRC1 complex and TMR-labeled the HALO-tagged CBX subunit of the full canonical PRC1 complex. For differently colored arrays, we used the G5E4 arrays labeled with one ATTO-647N dye and two biotins as well as G5E4 arrays labeled with one Cy3N dye and two biotins. After the 30-minute incubation of the array/PRC1 mixtures, we mixed the solutions of the differently colored arrays or PRC1 (depending on the experiment) and incubated these mixtures for an additional 15 minutes at 30°C. Afterwards, we prepared the flow-cell with bilayers as described above and added the mixtures to the flow-cell for imaging.

For the PRC1 redistribution experiments shown in **Figure S6H-J**, we first let condensates of ATTO-647N labeled nucleosomal arrays and mGFP-tagged PRC1 form for 30 minutes at 30°C. Afterwards, we added Cy3N labeled nucleosomal arrays at various concentrations and incubated for another 15 minutes at 30°C.

For all experiments, we note that the ratio of ATP, magnesium, and EDTA is critical. A very small change in concentrations can lead to different results (e.g. condensation of arrays alone when there is too much magnesium).

### Preparation of flow-cells for epifluorescence and confocal microscopy (droplet assays)

We typically mixed nucleosomal arrays (30 nM nucleosome final concentration unless otherwise noted) and various concentrations of PRC1 in TIRF Buffer (13 mM HEPES at pH 7.9, 120 mM KCl, 1.4 mM MgCl_2_, 0.25 mM EDTA, 0.9 mM ATP, 1% glycerol, 0.04 μg/μl BSA, 0.4 μg/μl kappa casein). For all experiments, we used G5E4 arrays labeled with either one Cy3N dye (for colocalization experiment with differently colored arrays and for FRAP data collection) or one ATTO-647N dye (for all other experiments) and no biotin as described above. Note, that we used different KCl concentrations for some experiments. If we did, it is clearly noted in the relevant section of the manuscript. We then incubated the 30 μl reaction at 30°C for 30 minutes in a thermocycler. In the meantime, we prepared the flow-cells by washing twice with TIRF buffer. After incubation, we added the samples to the slide and incubated for 5 minutes before imaging. For the experiments with PHC1 or PHC2 which have a C-terminal TEV site and a HALO-tag^49^, we added 0.01 μg/μl TEV protease.

For the mixing and colocalization experiments of PRC1 and arrays in different colors, we mixed PRC1 and nucleosomal arrays as described above. After the 30-minute incubation of the array/PRC1 mixtures, we mixed the solutions of the differently colored arrays or PRC1 (depending on the experiment) and incubated these mixtures for an additional 1, 15, 30, or 60 minutes at 30°C before adding them to the flow-cell for imaging.

### Preparation of flow-cells with pegylated glass

The flow-cells with mPEG-Succinimidyl Valerate, MW 5,000 (Laysan Bio., cat. no. mPEG-succinimidyl valerate, MW 5,000) and Biotin-PEG-SVA, MW 5,000 (Laysan Bio., cat. no. Biotin-PEG-SVA, MW 5,000) at a ratio of 80:1 were pegylated as previously described^78,79^. Once the coverslips (Zeiss, 474030-9000-000) and 24×50 mm microscope slides (Thermo Fisher Scientific, 50-311-35) were pegylated they were assembled with double-sided tape as described above. Samples were applied to pegylated flow-cells and TIRF imaged as described above for the bilayer system.

### Cell culture and generation of cell lines for doxycycline-inducible expression of mEGFP-CBX2 and mEGFP-CBX7

NIH-3T3 fibroblasts (American Type Culture Collection [ATCC]) were cultured in DMEM (Thermo Fisher Scientific, 11965092) supplemented with fetal calf serum to 10% (v/v) concentration and 1% penicillin/streptomycin (v/v). For the mEGFP-CBX2 experiments we used the previously published stable cell line^19^. The mEGFP-CBX7 cDNA was cloned into a pTRIPZ vector (Dharmacon). Subsequently HEK293T cells were transfected with this vector alongside second generation lentiviral packaging plasmids, pCMV-dR8.91 containing gag, pol, and rev genes and pMD2.G encoding the VSV-G envelope protein using the TransIT-Lenti transfection reagent (Mirus, MIR 6650). The virus was harvested after 48 hours and filtered through a 0.45 μm filter. Then, NIH-3T3 fibroblasts were transduced with lentivirus. After 48 hours, the cells were selected with 1 μg/ml puromycin.

We cloned the cDNA of mScarlet-PHC2 and mScarlet-PHC2 L307R into pEF1alpha vectors which were a kind gift from Constance L. Cepko (Addgene, Plasmid #11154)^80^. Cells for live cell imaging were plated at 0.2 x 10^4^ cells per ml in Poly-L-Lysine coated 8-well μ-Slide chambers (Ibidi, 80804). The next day, stable 3T3 cell lines for doxycycline-inducible expression of mEGFP-CBX2 or mEGFP-CBX7 were transiently transfected with either 200 ng of mScarlet-PHC2 or mScarlet-PHC2 L307R DNA using Lipofectamine LTX (Thermo Fisher Scientific, 15338030) according to manufacturer’s instructions. At the same time, we added doxycycline (1 μg/ml) in OptiMEM (Thermo Fisher Scientific, 31985062). After 24 hours, we changed the media to FluoroBrite (Thermo Fisher Scientific, A1896701) imaging media supplemented with fetal calf serum to 10% (v/v) concentration, 1% penicillin/streptomycin (v/v), and 20 mM HEPES pH 7.4.

### TIRF and epi-fluorescence microscopy data collection

The TIRF and epi-fluorescence data collections were conducted using a Nikon Ti-E inverted microscope equipped with a Ti-ND6-PFS perfect focus system and an APO TIRF 100x oil/1.49 DIC objective (Nikon). The microscope was also equipped with a Nikon-encoded x-y motorized stage and a piezo z-stage, an EMCCD camera (Andor iXon Ultra, DU-897U-CSO-#BV), and 488 nm, 561 nm and 640 nm laser. All imaging was performed with subsequent exposures of 102 msec, using the CCD mode (i.e. no EM gain) with 3 MHz and 16-bit. For all imaging, except for colocalization experiments we used a Quad filter cube 405/488/561/640nm (Chroma, TRF89901). For colocalization experiments, we used cubes specifically for each laser line to minimize crosstalk (Chroma, TRF49904 (488 nm), TRF49909 (561 nm), TRF49914 (640 nm)). The movies for the dynamic studies of condensate formation in **Figure S6A-E** were acquired with 20 second intervals.

### FRAP data collection

FRAP data was collected on a Nikon A1R laser-scanning confocal inverted microscope equipped with a 60× oil immersion objective. We collected data with a scanning speed of 1/2 frames / second over an area of 1024×1024 pixels and with 2× averaging. We selected “channel series” to avoid crosstalk, set the pinhole to 0.8 and used a zoom of 3.0 resulting in a 0.07 μm pixel size. The 488 nm laser was set to HV=80, Offset=0, and Power=1.00. The 561 nm laser was set to HV=100, Offset=0, and Power=10.00. For bleaching we set the laser power to 100% for the respective laser line. The data was collected as follows. We first imaged 3 loops with a 45 second interval before bleaching, then bleached over 15 loops with no delay (i.e. ∼ 15 seconds), and collected 40 more loops with 45 second intervals.

### Fluorescent microscopy of 3T3 fibroblasts

Images of 3T3 fibroblasts were acquired on a Nikon A1R laser-scanning confocal inverted microscope equipped with a 60× oil immersion objective. We collected data with a scanning speed of 1/16 frames / second over an area of 1024×1024 pixels and with 2× averaging. We selected “channel series” to avoid crosstalk, set the pinhole to 0.8 and used a zoom of 3.0 resulting in a 0.07 μm pixel size. The 488 nm laser was set to HV=140, Offset=-30, and Power=1.00. The 561 nm laser was set to HV=140, Offset=-20, and Power=2.00. For all imaging 5.0% of CO_2_ was supplied and a temperature of 37°C was maintained.

### TIRF and epi-fluorescence microscopy data analysis

The raw microscopy files were analyzed in Fiji (light microscopy data) and Python (version 3.9.6, Python Software Foundation). To extract values such as the intensity for each spot for all the images, we wrote a custom Fiji macro. Briefly, we first opened the images with Bio-Formats and created a mask for all spots. To this end, the images were converted to an 8-bit file, the background was subtracted, and an auto threshold (intermodes dark) was applied. Afterwards, we applied the Watershed algorithm to separate spots that might appear as one but are actually two separate ones. With this mask, we then measured the raw intensity values of each spot and exported these values as .txt files. In addition, these .txt files also contained the area and perimeter of each spot; values which are required to calculate the roundness per spot. We also calculated the intensity of the entire image minus the areas where we detected spots to calculate the background intensity and exported this as .txt files. These .txt files were then further processed in a custom python script, in which the background intensity was divided into 36 equal squares to better determine the local background per area. We then subtracted the background intensity from the intensities that we measured for each spot. This background subtracted intensity values were finally used to plot various graphs presented in this manuscript.

To convert the intensity values of each spot to the number of molecules, we used beaching traces and calculated the average number of photons per nucleosomal array or per PRC1 molecule. We typically collected bleaching traces for a mixture of nucleosomal arrays with 5 nM nucleosomes and 2.5 nM PRC1 so that we did not get condensates and only had a few bleaching steps per spot. To determine the bleaching steps we used quickPBSA^81^.

To calculate the percentage of molecules that are found in condensates we set the following cut-offs: every spot with more than 350 PRC1 molecules or more than 20 nucleosomal arrays (i.e. 240 nucleosomes for a 12mer nucleosomal array) was considered a condensate. If we changed the cut-off value, the sigmoidal curve was shifted, but the overall shape of the curve did not change indicating that our observations held true independent of the chosen cut-off values. We note that we only used spots that were in the field of view for our calculation. This means that anything that was in solution and not captured with our TIRF imaging, was not taken into consideration. However, we found that typically the small condensates or single nucleosomal arrays bound to the lipid bilayer before the larger condensates and thus predict that we potentially missed more larger condensates than single nucleosomal arrays and thus, if anything underestimated the number of molecules in condensates.

To calculate the correlation for the mixing experiments of differently colored PRC1, we created a mask using the intensity of the nucleosomal arrays (**Figure 2**). The mask for nucleosomal arrays was created as described above. We then used this mask and measured the background subtracted intensity for each of the two differently colored PRC1s at each pixel of the mask. For the differently colored nucleosomal arrays, we created a mask based on the PRC1 intensity and proceeded in the same way as for differently colored PRC1 molecules.

### FRAP data analysis

As with the TIRF and epi-fluorescence microscopy data, we loaded the images into Fiji. Since we noticed drift for some of the movies, we corrected the drift by running StackReg^82^. Then, we created masks to measure the intensities as described above. However, here we distinguished between areas that were bleached and areas that were not bleached and created separate .txt files. The not bleached spots were used for a general photobleaching correction. We also normalized the intensity trace for each bleached spot so that it was at 1 before the bleaching occurred and dropped to 0 at the moment of bleaching. The mobile fraction was estimated as a percentage of fluorescence intensity that recovered until the last frame.

### Analysis of live cell imaging data

We first applied a deconvolution to all images using the “2D Deconvolution” in Nikon Elements with the “Fast” method and “Automatic” Noise Level setting. Afterwards, the data was loaded into Fiji. We used the “Spot intensity” plug-in (https://imagej.net/plugins/spot-intensity-analysis) to count the number of spots and to measure their intensities with the following settings: Time interval of 1.0 s, Electrons per ADU of 1.0, Check first n frames of 1, Spot radius of 4, Background of 2,000, and Background estimation Median. Subsequently, the data was further analyzed and plotted using a custom python script.

### Figure preparation

All figures and graphs were prepared by either using Fiji (light microscopy data), Affinity designer (version 1.6.1, Serif (Europe) Ltd), or Python (version 3.9.6, Python Software Foundation).

## References

1. Piunti, A., and Shilatifard, A. (2021). The roles of Polycomb repressive complexes in mammalian development and cancer. Nat. Rev. Mol. Cell Biol. 22, 326–345. 10.1038/s41580-021-00341-1.

2. Mehta, S., and Zhang, J. (2022). Liquid–liquid phase separation drives cellular function and dysfunction in cancer. Nat. Rev. Cancer 22, 239–252. 10.1038/s41568-022-00444-7.

3. Flavahan, W.A., Gaskell, E., and Bernstein, B.E. (2017). Epigenetic plasticity and the hallmarks of cancer. Science 357. 10.1126/science.aal2380.

4. Shen, H., and Laird, P.W. (2013). Interplay between the Cancer Genome and Epigenome. Cell 153, 38–55. 10.1016/j.cell.2013.03.008.

5. Hildebrand, E.M., and Dekker, J. (2020). Mechanisms and Functions of Chromosome Compartmentalization. Trends Biochem. Sci. 45, 385–396. 10.1016/j.tibs.2020.01.002.

6. Klemm, S.L., Shipony, Z., and Greenleaf, W.J. (2019). Chromatin accessibility and the regulatory epigenome. Nat. Rev. Genet. 20, 207–220. 10.1038/s41576-018-0089-8.

7. Kingston, R.E., and Tamkun, J.W. (2014). Transcriptional Regulation by Trithorax-Group Proteins. Cold Spring Harb. Perspect. Biol. 6, a019349–a019349. 10.1101/cshperspect.a019349.

8. Blackledge, N.P., and Klose, R.J. (2021). The molecular principles of gene regulation by Polycomb repressive complexes. Nat. Rev. Mol. Cell Biol. 22, 815–833. 10.1038/s41580-021-00398-y.

9. Kim, J.J. Context-specific Polycomb mechanisms in development. 16.

10. Schuettengruber, B., Bourbon, H.-M., Croce, L.D., and Cavalli, G. (2017). Genome Regulation by Polycomb and Trithorax: 70 Years and Counting. Cell 171, 34–57. 10.1016/j.cell.2017.08.002.

11. Kuroda, M.I., Kang, H., De, S., and Kassis, J.A. (2020). Dynamic Competition of Polycomb and Trithorax in Transcriptional Programming. Annu. Rev. Biochem. 89, annurev-biochem-120219-103641. 10.1146/annurev-biochem-120219-103641.

12. Aranda, S., Mas, G., and Croce, L.D. (2015). Regulation of gene transcription by Polycomb proteins. Sci. Adv. 1, e1500737. 10.1126/sciadv.1500737.

13. Kundu, S., Ji, F., Sunwoo, H., Jain, G., Lee, J.T., Sadreyev, R.I., Dekker, J., and Kingston, R.E. (2017). Polycomb Repressive Complex 1 Generates Discrete Compacted Domains that Change during Differentiation. Mol. Cell 65, 432–446.e5. 10.1016/j.molcel.2017.01.009.

14. Isono, K., Endo, T.A., Ku, M., Yamada, D., Suzuki, R., Sharif, J., Ishikura, T., Toyoda, T., Bernstein, B.E., and Koseki, H. (2013). SAM Domain Polymerization Links Subnuclear Clustering of PRC1 to Gene Silencing. Dev. Cell 26, 565–577. 10.1016/j.devcel.2013.08.016.

15. Wani, A.H., Boettiger, A.N., Schorderet, P., Ergun, A., Münger, C., Sadreyev, R.I., Zhuang, X., Kingston, R.E., and Francis, N.J. (2016). Chromatin topology is coupled to Polycomb group protein subnuclear organization. Nat. Commun. 7, 10291. 10.1038/ncomms10291.

16. Saurin, A.J., Shiels, C., Williamson, J., Satijn, D.P.E., Otte, A.P., Sheer, D., and Freemont, P.S. (1998). The Human Polycomb Group Complex Associates with Pericentromeric Heterochromatin to Form a Novel Nuclear Domain. J. Cell Biol. 142, 887–898. 10.1083/jcb.142.4.887.

17. Satijn, D.P.E., Gunster, M.J., van der Vlag, J., Hamer, K.M., Schul, W., Alkema, M.J., Saurin, A.J., Freemont, P.S., van Driel, R., and Otte, A.P. (1997). RING1 Is Associated with the Polycomb Group Protein Complex and Acts as a Transcriptional Repressor. Mol. Cell. Biol. 17, 4105–4113. 10.1128/MCB.17.7.4105.

18. Tatavosian, R., Kent, S., Brown, K., Yao, T., Duc, H.N., Huynh, T.N., Zhen, C.Y., Ma, B., Wang, H., and Ren, X. (2019). Nuclear condensates of the Polycomb protein chromobox 2 (CBX2) assemble through phase separation. J. Biol. Chem. 294, 1451–1463. 10.1074/jbc.RA118.006620.

19. Plys, A.J., Davis, C.P., Kim, J., Rizki, G., Keenen, M.M., Marr, S.K., and Kingston, R.E. (2019). Phase separation of Polycomb-repressive complex 1 is governed by a charged disordered region of CBX2. Genes Dev. 33, 799–813. 10.1101/gad.326488.119.

20. Sabari, B.R., Dall’Agnese, A., and Young, R.A. (2020). Biomolecular Condensates in the Nucleus. Trends Biochem. Sci. 45, 961–977. 10.1016/j.tibs.2020.06.007.

21. Lyon, A.S., Peeples, W.B., and Rosen, M.K. (2021). A framework for understanding the functions of biomolecular condensates across scales. Nat. Rev. Mol. Cell Biol. 22, 215–235. 10.1038/s41580-020-00303-z.

22. Narlikar, G.J. (2020). Phase-separation in chromatin organization. J. Biosci. 45, 5. 10.1007/s12038-019-9978-z.

23. Lafontaine, D.L.J., Riback, J.A., Bascetin, R., and Brangwynne, C.P. (2021). The nucleolus as a multiphase liquid condensate. Nat. Rev. Mol. Cell Biol. 22, 165–182. 10.1038/s41580-020-0272-6.

24. Rippe, K. (2022). Liquid–Liquid Phase Separation in Chromatin. Cold Spring Harb. Perspect. Biol. 14, a040683. 10.1101/cshperspect.a040683.

25. Gibson, B.A., Doolittle, L.K., Schneider, M.W.G., Jensen, L.E., Gamarra, N., Henry, L., Gerlich, D.W., Redding, S., and Rosen, M.K. (2019). Organization of Chromatin by Intrinsic and Regulated Phase Separation. Cell 179, 470–484.e21. 10.1016/j.cell.2019.08.037.

26. Jain, A., and Vale, R.D. (2017). RNA phase transitions in repeat expansion disorders. Nature 546, 243–247. 10.1038/nature22386.

27. Sabari, B.R., Dall’Agnese, A., Boija, A., Klein, I.A., Coffey, E.L., Shrinivas, K., Abraham, B.J., Hannett, N.M., Zamudio, A.V., Manteiga, J.C., et al. (2018). Coactivator condensation at super-enhancers links phase separation and gene control. Science 361, eaar3958. 10.1126/science.aar3958.

28. Shrinivas, K., Sabari, B.R., Coffey, E.L., Klein, I.A., Boija, A., Zamudio, A.V., Schuijers, J., Hannett, N.M., Sharp, P.A., Young, R.A., et al. (2019). Enhancer Features that Drive Formation of Transcriptional Condensates. Mol. Cell 75, 549–561.e7. 10.1016/j.molcel.2019.07.009.

29. Guo, Y.E., Manteiga, J.C., Henninger, J.E., Sabari, B.R., Dall’Agnese, A., Hannett, N.M., Spille, J.-H., Afeyan, L.K., Zamudio, A.V., Shrinivas, K., et al. (2019). Pol II phosphorylation regulates a switch between transcriptional and splicing condensates. Nature 572, 543–548. 10.1038/s41586-019-1464-0.

30. Cho, W.-K., Spille, J.-H., Hecht, M., Lee, C., Li, C., Grube, V., and Cisse, I.I. (2018). Mediator and RNA polymerase II clusters associate in transcription-dependent condensates. Science 361, 412–415. 10.1126/science.aar4199.

31. Gallego, L.D., Schneider, M., Mittal, C., Romanauska, A., Gudino Carrillo,R.M., Schubert, T., Pugh, B.F., and Köhler, A. (2020). Phase separation directs ubiquitination of gene-body nucleosomes. Nature 579, 592–597. 10.1038/s41586-020-2097-z.

32. Lu, H., Yu, D., Hansen, A.S., Ganguly, S., Liu, R., Heckert, A., Darzacq, X., and Zhou, Q. (2018). Phase-separation mechanism for C-terminal hyperphosphorylation of RNA polymerase II. Nature 558, 318–323. 10.1038/s41586-018-0174-3.

33. Boehning, M., Dugast-Darzacq, C., Rankovic, M., Hansen, A.S., Yu, T., Marie-Nelly, H., McSwiggen, D.T., Kokic, G., Dailey, G.M., Cramer, P., et al. (2018). RNA polymerase II clustering through carboxy-terminal domain phase separation. Nat. Struct. Mol. Biol. 25, 833–840. 10.1038/s41594-018-0112-y.

34. Guo, C., Luo, Z., and Lin, C. (2021). Phase separation, transcriptional elongation control, and human diseases. J. Mol. Cell Biol. 13, 314–318. 10.1093/jmcb/mjab023.

35. Boija, A., Klein, I.A., Sabari, B.R., Dall’Agnese, A., Coffey, E.L., Zamudio, A.V., Li, C.H., Shrinivas, K., Manteiga, J.C., Hannett, N.M., et al. (2018). Transcription Factors Activate Genes through the Phase-Separation Capacity of Their Activation Domains. Cell 175, 1842–1855.e16. 10.1016/j.cell.2018.10.042.

36. Larson, A.G., Elnatan, D., Keenen, M.M., Trnka, M.J., Johnston, J.B., Burlingame, A.L., Agard, D.A., Redding, S., and Narlikar, G.J. (2017). Liquid droplet formation by HP1α suggests a role for phase separation in heterochromatin. Nature 547, 236–240. 10.1038/nature22822.

37. Strom, A.R., Biggs, R.J., Banigan, E.J., Wang, X., Chiu, K., Herman, C., Collado, J., Yue, F., Ritland Politz, J.C., Tait, L.J., et al. (2021). HP1α is a chromatin crosslinker that controls nuclear and mitotic chromosome mechanics. eLife 10, e63972. 10.7554/eLife.63972.

38. Seif, E., Kang, J.J., Sasseville, C., Senkovich, O., Kaltashov, A., Boulier, E.L., Kapur, I., Kim, C.A., and Francis, N.J. (2020). Phase separation by the polyhomeotic sterile alpha motif compartmentalizes Polycomb Group proteins and enhances their activity. Nat. Commun. 11, 5609. 10.1038/s41467-020-19435-z.

39. Hansen, J.C., Maeshima, K., and Hendzel, M.J. (2021). The solid and liquid states of chromatin. Epigenetics Chromatin 14, 50. 10.1186/s13072-021-00424-5.

40. Wang, L., Gao, Y., Zheng, X., Liu, C., Dong, S., Li, R., Zhang, G., Wei, Y., Qu, H., Li, Y., et al. (2019). Histone Modifications Regulate Chromatin Compartmentalization by Contributing to a Phase Separation Mechanism. Mol. Cell 76, 646–659.e6. 10.1016/j.molcel.2019.08.019.

41. Morey, L., Pascual, G., Cozzuto, L., Roma, G., Wutz, A., Benitah, S.A., and Di Croce, L. (2012). Nonoverlapping Functions of the Polycomb Group Cbx Family of Proteins in Embryonic Stem Cells. Cell Stem Cell 10, 47–62. 10.1016/j.stem.2011.12.006.

42. Kim, J., and Kingston, R.E. (2020). The CBX family of proteins in transcriptional repression and memory. J. Biosci. 45, 16. 10.1007/s12038-019-9972-5.

43. Grau, D.J., Chapman, B.A., Garlick, J.D., Borowsky, M., Francis, N.J., and Kingston, R.E. (2011). Compaction of chromatin by diverse Polycomb group proteins requires localized regions of high charge. Genes Dev. 25, 2210–2221. 10.1101/gad.17288211.

44. Lau, M.S., Schwartz, M.G., Kundu, S., Savol, A.J., Wang, P.I., Marr, S.K., Grau, D.J., Schorderet, P., Sadreyev, R.I., Tabin, C.J., et al. (2017). Mutation of a nucleosome compaction region disrupts Polycomb-mediated axial patterning. Science 355, 1081–1084. 10.1126/science.aah5403.

45. Coré, N., Bel, S., Gaunt, S.J., Aurrand-Lions, M., Pearce, J., Fisher, A., and Djabali, M. Altered cellular proliferation and mesoderm patterning in Polycomb-M33deficient mice.

46. Kim, C.A., Gingery, M., Pilpa, R.M., and Bowie, J.U. (2002). The SAM domain of polyhomeotic forms a helical polymer. Nat. Struct. Biol. 9, 5.

47. Flaus, A. (2011). Principles and practice of nucleosome positioning in vitro. Front. Life Sci. 5, 5–27. 10.1080/21553769.2012.702667.

48. Utley, R.T., Ikeda, K., Grant, P.A., Côté, J., Steger, D.J., Eberharter, A., John, S., and Workman, J.L. (1998). Transcriptional activators direct histone acetyltransferase complexes to nucleosomes. Nature 394, 498–502. 10.1038/28886.

49. Los, G.V., Encell, L.P., McDougall, M.G., Hartzell, D.D., Karassina, N., Zimprich, C., Wood, M.G., Learish, R., Ohana, R.F., Urh, M., et al. (2008). HaloTag: A Novel Protein Labeling Technology for Cell Imaging and Protein Analysis. ACS Chem. Biol. 3, 373–382. 10.1021/cb800025k.

50. Jaensch, E.S., Zhu, J., Cochrane, J.C., Marr, S.K., Oei, T.A., Damle, M., McCaslin, E.Z., and Kingston, R.E. (2021). A Polycomb domain found in committed cells impairs differentiation when introduced into PRC1 in pluripotent cells. Mol. Cell 81, 4677–4691.e8. 10.1016/j.molcel.2021.09.018.

51. Kapur, I., Boulier, E.L., and Francis, N.J. (2022). Regulation of Polyhomeotic Condensates by Intrinsically Disordered Sequences That Affect Chromatin Binding. Epigenomes 6, 40. 10.3390/epigenomes6040040.

52. Gambetta, M.C., and Müller, J. (2014). O-GlcNAcylation Prevents Aggregation of the Polycomb Group Repressor Polyhomeotic. Dev. Cell 31, 629–639. 10.1016/j.devcel.2014.10.020.

53. Morin, J.A., Wittmann, S., Choubey, S., Klosin, A., Golfier, S., Hyman, A.A., Jülicher, F., and Grill, S.W. (2022). Sequence-dependent surface condensation of a pioneer transcription factor on DNA. Nat. Phys. 18, 271–276. 10.1038/s41567-021-01462-2.

54. Spakman, D., King, G.A., Peterman, E.J.G., and Wuite, G.J.L. (2020). Constructing arrays of nucleosome positioning sequences using Gibson Assembly for single-molecule studies. Sci. Rep. 10, 9903. 10.1038/s41598-020-66259-4.

55. Banani, S.F., Rice, A.M., Peeples, W.B., Lin, Y., Jain, S., Parker, R., and Rosen, M.K. (2016). Compositional Control of Phase-Separated Cellular Bodies. Cell 166, 651–663. 10.1016/j.cell.2016.06.010.

56. Keenen, M.M., Brown, D., Brennan, L.D., Renger, R., Khoo, H., Carlson, C.R., Huang, B., Grill, S.W., Narlikar, G.J., and Redding, S. (2021). HP1 proteins compact DNA into mechanically and positionally stable phase separated domains. eLife 10, e64563. 10.7554/eLife.64563.

57. Frank, L., and Rippe, K. (2020). Repetitive RNAs as Regulators of Chromatin-Associated Subcompartment Formation by Phase Separation. J. Mol. Biol. 432, 4270–4286. 10.1016/j.jmb.2020.04.015.

58. Jack, A., Kim, Y., Strom, A.R., Lee, D.S.W., Williams, B., Schaub, J.M., Kellogg, E.H., Finkelstein, I.J., Ferro, L.S., Yildiz, A., et al. (2022). Compartmentalization of telomeres through DNA-scaffolded phase separation. Dev. Cell 57, 277–290.e9. 10.1016/j.devcel.2021.12.017.

59. Fan, C., Zhang, H., Fu, L., Li, Y., Du, Y., Qiu, Z., and Lu, F. (2020). Rett mutations attenuate phase separation of MeCP2. Cell Discov. 6, 38. 10.1038/s41421-020-0172-0.

60. Li, C.H., Coffey, E.L., Dall’Agnese, A., Hannett, N.M., Tang, X., Henninger, J.E., Platt, J.M., Oksuz, O., Zamudio, A.V., Afeyan, L.K., et al. (2020). MeCP2 links heterochromatin condensates and neurodevelopmental disease. Nature 586, 440–444. 10.1038/s41586-020-2574-4.

61. Wang, L., Hu, M., Zuo, M.-Q., Zhao, J., Wu, D., Huang, L., Wen, Y., Li, Y., Chen, P., Bao, X., et al. (2020). Rett syndrome-causing mutations compromise MeCP2-mediated liquid–liquid phase separation of chromatin. Cell Res. 30, 393–407. 10.1038/s41422-020-0288-7.

62. Uckelmann, M., Levina, V., Taveneau, C., Ng, X.H., Pandey, V., Zhang, Q., Flanigan, S., Li, M., Das, P.P., Marco, A. de, et al. (2023). Dynamic PRC1-CBX8 stabilizes a porous structure of chromatin condensates. 2023.05.08.539931. 10.1101/2023.05.08.539931.

63. Huseyin, M.K., and Klose, R.J. (2021). Live-cell single particle tracking of PRC1 reveals a highly dynamic system with low target site occupancy. Nat. Commun. 12, 887. 10.1038/s41467-021-21130-6.

64. Zhen, C.Y., Tatavosian, R., Huynh, T.N., Duc, H.N., Das, R., Kokotovic, M., Grimm, J.B., Lavis, L.D., Lee, J., Mejia, F.J., et al. (2016). Live-cell single-molecule tracking reveals co-recognition of H3K27me3 and DNA targets polycomb Cbx7-PRC1 to chromatin. eLife 5, e17667. 10.7554/eLife.17667.

65. Gambetta, M.C., Oktaba, K., and Müller, J. (2009). Essential Role of the Glycosyltransferase Sxc/Ogt in Polycomb Repression. Science 325, 93–96. 10.1126/science.1169727.

66. Sinclair, D.A.R., Syrzycka, M., Macauley, M.S., Rastgardani, T., Komljenovic, I., Vocadlo, D.J., Brock, H.W., and Honda, B.M. (2009). Drosophila O-GlcNAc transferase (OGT) is encoded by the Polycomb group (PcG) gene, super sex combs (sxc). Proc. Natl. Acad. Sci. 106, 13427–13432. 10.1073/pnas.0904638106.

67. Kawaguchi, T., Machida, S., Kurumizaka, H., Tagami, H., and Nakayama, J. (2017). Phosphorylation of CBX2 controls its nucleosome-binding specificity. J. Biochem. (Tokyo) 162, 343–355. 10.1093/jb/mvx040.

68. Lavigne, M., Francis, N.J., King, I.F.G., and Kingston, R.E. (2004). Propagation of Silencing: Recruitment and Repression of Naive Chromatin In trans by Polycomb Repressed Chromatin. Mol. Cell 13, 415–425. 10.1016/S1097-2765(04)00006-1.

69. Brown, K., Chew, P.Y., Ingersoll, S., Espinosa, J.R., Aguirre, A., Kutateladze, T., Guevara, R.C., and Ren, X. (2022). Principles of assembly and regulation of condensates of Polycomb repressive complex 1 through phase separation (Biochemistry) 10.1101/2022.12.26.521954.

70. Weissmann, F., Petzold, G., VanderLinden, R., Huis in ‘t Veld, P.J., Brown, N.G., Lampert, F., Westermann, S., Stark, H., Schulman, B.A., and Peters, J.-M. (2016). biGBac enables rapid gene assembly for the expression of large multisubunit protein complexes. Proc. Natl. Acad. Sci. 113, E2564–E2569. 10.1073/pnas.1604935113.

71. Dyer, P.N., Edayathumangalam, R.S., White, C.L., Bao, Y., Chakravarthy, S., Muthurajan, U.M., and Luger, K. (2003). Reconstitution of Nucleosome Core Particles from Recombinant Histones and DNA. In Methods in Enzymology (Elsevier), pp.23–44. 10.1016/S0076-6879(03)75002-2.

72. Neumann, H., Hancock, S.M., Buning, R., Routh, A., Chapman, L., Somers, J., Owen-Hughes, T., van Noort, J., Rhodes, D., and Chin, J.W. (2009). A Method for Genetically Installing Site-Specific Acetylation in Recombinant Histones Defines the Effects of H3 K56 Acetylation. Mol. Cell 36, 153–163. 10.1016/j.molcel.2009.07.027.

73. Luger, K., Rechsteiner, T.J., and Richmond, T.J. (1999). Preparation of nucleosome core particle from recombinant histones. In Methods in Enzymology (Elsevier), pp.3–19. 10.1016/S0076-6879(99)04003-3.

74. Nair, P.M., Salaita, K., Petit, R.S., and Groves, J.T. (2011). Using patterned supported lipid membranes to investigate the role of receptor organization in intercellular signaling. Nat. Protoc. 6, 523–539. 10.1038/nprot.2011.302.

75. Qi, Z., and Greene, E.C. (2016). Visualizing recombination intermediates with single-stranded DNA curtains. Methods 105, 62–74. 10.1016/j.ymeth.2016.03.027.

76. Ma, C.J., Steinfeld, J.B., and Greene, E.C. (2017). Chapter Eight Single-Stranded DNA Curtains for Studying Homologous Recombination. In Methods in Enzymology Single-Molecule Enzymology: Nanomechanical Manipulation and Hybrid Methods., M. Spies and Y. R. Chemla, eds. (Academic Press), pp.193–219. 10.1016/bs.mie.2016.08.005.

77. Niekamp, S., Stuurman, N., and Vale, R.D. (2020). A 6-nm ultra-photostable DNA FluoroCube for fluorescence imaging. Nat. Methods 17, 437–441. 10.1038/s41592-020-0782-3.

78. Jain, A., Liu, R., Xiang, Y.K., and Ha, T. (2012). Single-molecule pull-down for studying protein interactions. Nat. Protoc. 7, 445–452. 10.1038/nprot.2011.452.

79. Haque, F., Freniere, C., Ye, Q., Mani, N., Wilson-Kubalek, E.M., Ku, P.-I., Milligan, R.A., and Subramanian, R. (2022). Cytoskeletal regulation of a transcription factor by DNA mimicry via coiled-coil interactions. Nat. Cell Biol. 24, 1088–1098. 10.1038/s41556-022-00935-7.

80. Matsuda, T., and Cepko, C.L. (2004). Electroporation and RNA interference in the rodent retina in vivo and in vitro. Proc. Natl. Acad. Sci. 101, 16–22. 10.1073/pnas.2235688100.

81. Hummert, J., Yserentant, K., Fink, T., Euchner, J., Ho, Y.X., Tashev, S.A., and Herten, D.-P. (2021). Photobleaching step analysis for robust determination of protein complex stoichiometries. Mol. Biol. Cell 32, ar35. 10.1091/mbc.E20-09-0568.

82. Thevenaz, P., Ruttimann, U.E., and Unser, M. (1998). A pyramid approach to subpixel registration based on intensity. IEEE Trans. Image Process. 7, 27–41. 10.1109/83.650848.

