## Supplementary-Information for "Modularity of PRC1 Composition and Chromatin Interaction define Condensate Properties"

### Supplementary Figures

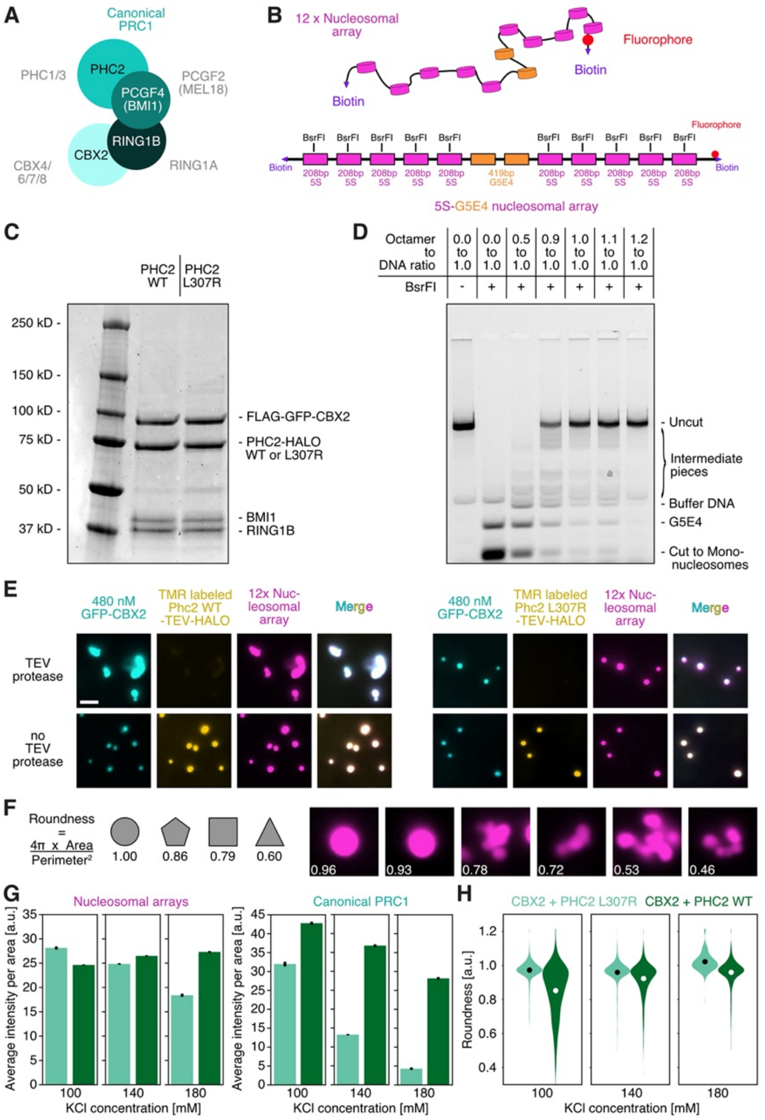

**Figure S1 | Design and assessment of canonical PRC1 and nucleosomal arrays.** (**A**) Schematic of canonical complex composition with its various paralogs listed in gray[1,2]. (**B**) Schematics of 12mer G5E4 nucleosomal arrays[3,4] which has been used for most of the experiments in this study. (**C**) 4-20% PAGE of purified canonical PRC1 complex with PHC2-TEV-HALO WT and PHC2-TEV-HALO L307R. (**D**) 1% Agarose gel to assess the extent of nucleosome occupancy of assembled nucleosomal arrays for various ratios of Octamer to DNA. Buffer DNA is added to prevent precipitation of unincorporated histones. (**E**) Micrographs of PRC1 complex mixed with nucleosomal arrays with or without TEV protease. Nucleosome concentration was 20 nM and PRC1 concentration was 480 nM. The scale bar is 5 µm. The data is representative of three technical repeats. (**F**) Definition of roundness criteria and example micrographs with roundness values. (**G**) Bar plot of average intensity per area for nucleosomal arrays (left) and PRC1 (right). PRC1 complex with mGFP-tagged CBX2 and either PHC2 WT (dark green) or PHC2

L307R (light green). Nucleosome concentration was 30 nM and PRC1 concentration was 610 nM. Error bar shows standard error of the mean. (**H**) Violin plot of PRC1 roundness of data as shown in **Figure 1F**. PRC1 complex with mGFP-tagged CBX2 and either PHC2 WT (dark green) or PHC2 L307R (light green). Circles represent the mean average value. (**G, H**) For each condition 325 to >1000 spots have been analyzed. Data are representative of at least three technical repeats.

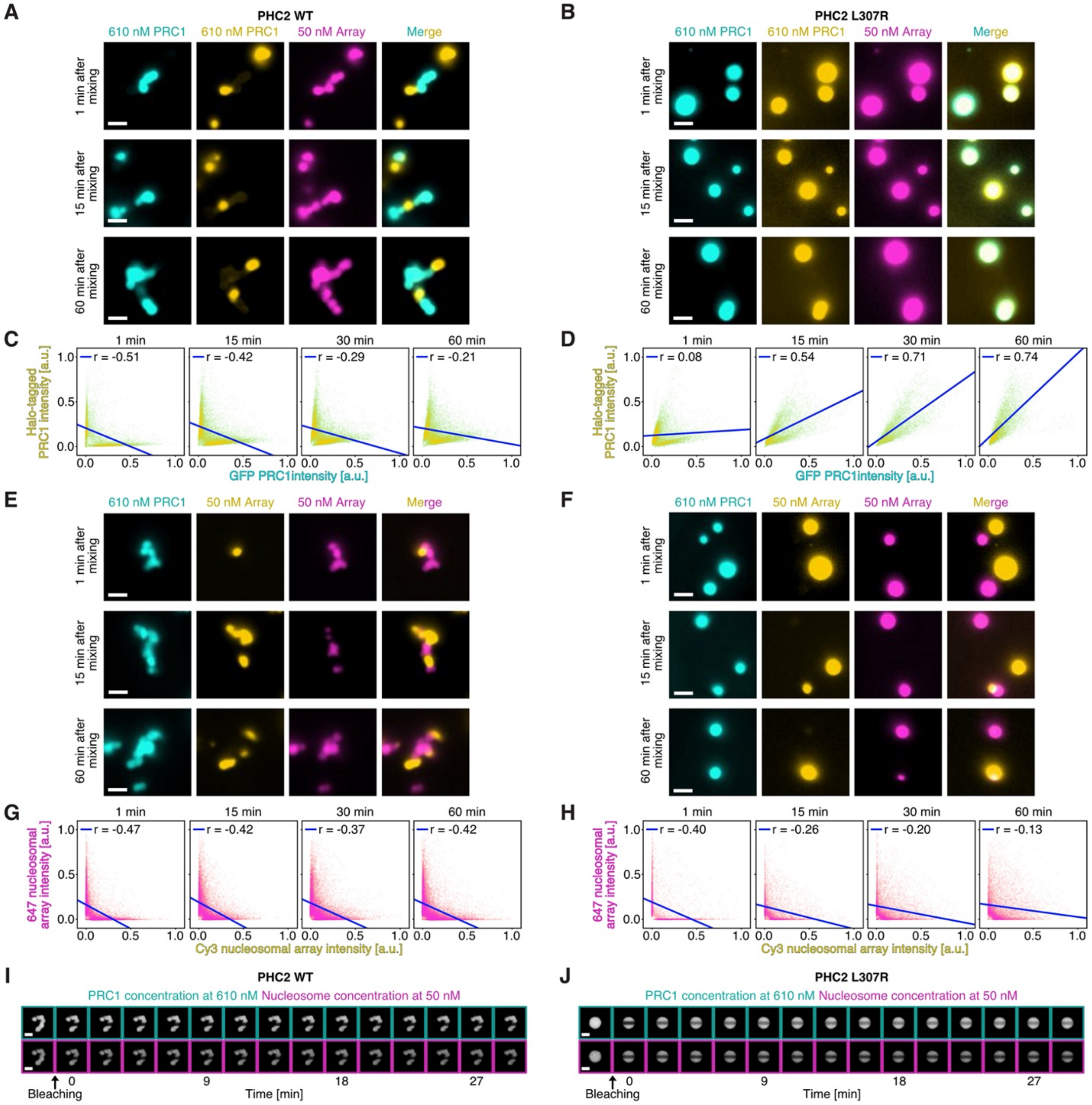


**Figure S2 | Effects of SAM polymerization on PRC1 and array dynamics in condensates.** (**A, B**) Representative micrographs of condensates with differently colored PRC1 molecules 1, 15, and 60 minutes after mixing for PRC1 complexes with (**A**) PHC2 WT and (**B**) PHC2 L307R. The scale bar is 5 µm. (**C, D**) Scatter plots for differently colored PRC1 with (**C**) PHC2 WT and (**D**) PHC2 L307R after 1, 15, 30, and 60 minutes of mixing condensates - related to **Figure 2**. The r values are the Pearson correlation coefficient. (**E, F**) Representative micrographs of condensates with differently colored nucleosomal arrays 1, 15, and 60 minutes after mixing for PRC1 complexes with (**E**) PHC2 WT and (**F**) PHC2 L307R. The scale bar is 5 µm. (**G, H**) Scatter plots for differently colored nucleosomal arrays with (**G**) PHC2 WT and (**H**) PHC2 L307R after 1, 15, 30, and 60 minutes of mixing condensates - related to **Figure 2.** The r values are the Pearson correlation coefficient. (**A-H**) Nucleosome concentration was 30 nM and PRC1 concentration was 610 nM. For each condition >100 spots have been analyzed. All data is representative of at least three technical repeats. (**I, J**) Images of nucleosomal arrays (magenta box) and canonical PRC1 (cyan box) with (**I**) PHC2 WT and (**J**) PHC2 L307R before and after bleaching - related to **Figure 2**. The scale bar is 2 μm.

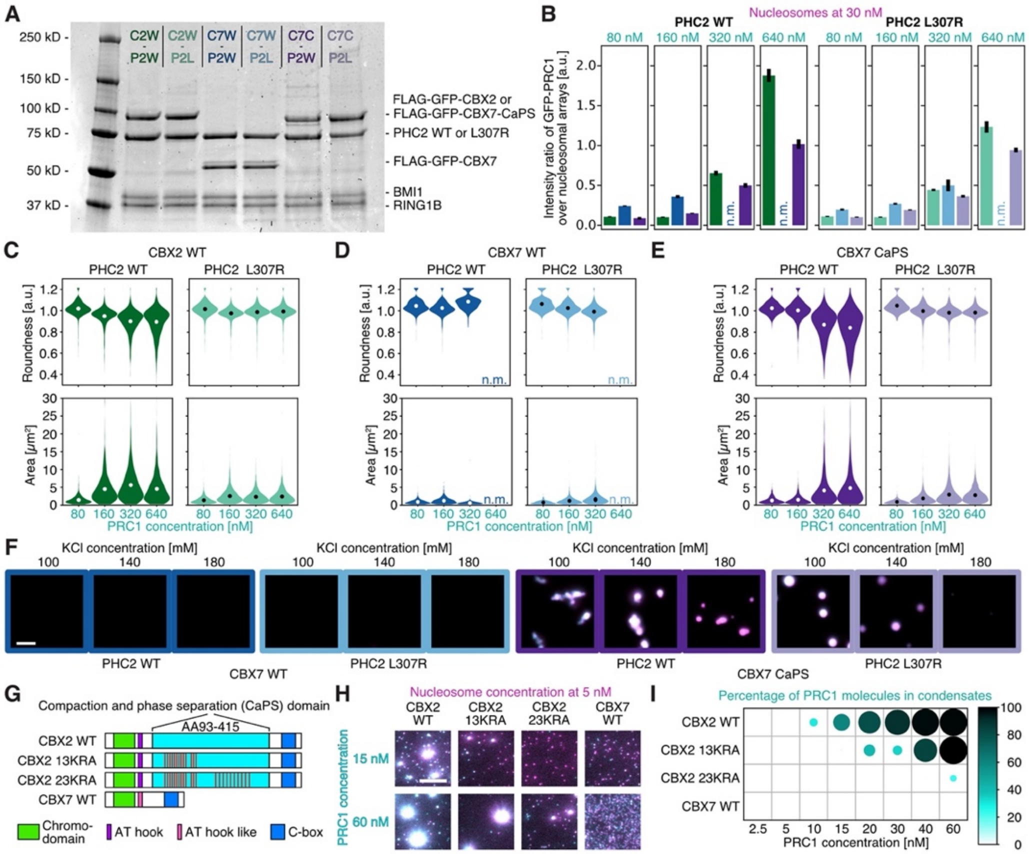


**Figure S3 | Contributions of CBX and PHC subunits to condensate formation and morphology.** (**A**) 4-20% SDS-PAGE of purified canonical PRC1 complexes with different CBX and PHC subunits. C2W, CBX2 WT. C7W, CBX7 WT. C7C, CBX7 CaPS. P2W, PHC2 WT. P2L, PHC2 L307R. Note that C2W-P2W and C2W-P2L are identical to **Figure S1C**. (**B**) Bar plot of intensity ratio of canonical PRC1 over nucleosomal arrays. Error bar is standard error of the mean. (**C-E**) Violin plots of puncta roundness and area for PRC1 complexes with (**C**) CBX2 WT, (**D**) CBX7 WT, and (**E**) CBX7 CaPS. The nucleosome concentration was 30 nM. For each condition >100 spots have been analyzed. All data is representative of at least three technical repeats. Circles represent the mean average value. N.m. is not measurable. (**F**) Example micrographs of nucleosomal arrays (magenta) and canonical PRC1 complex (cyan) with mGFP-tagged CBX7 and either PHC2 WT (dark blue box) or PHC2 L307R (light blue box) as well as with mGFP-tagged CBX7 CaPS and either PHC2 WT (dark violet box) or PHC2 L307R (light violet box). Nucleosome concentration was 30 nM and PRC1 concentration was 610 nM. The scale bar is 10 µm. (**B-E**) For each condition 325 to >1000 spots have been analyzed. Data are representative of at least three technical repeats. N.m. is not measurable. (**G**) Schematic of CBX subunits. For 13KRA and 23KRA, 13 or 23 arginine and lysine residues have been mutated to alanines[5–7]. (**H**) Representative TIRF micrographs of canonical PRC1 complex with PHC2 L307R and various CBX subunits as shown in **G**. The scale bar is 20 µm. (**I**) Percentage of PRC1 molecules in condensates for PRC1 complexes with various CBX subunits as shown in **G**. Circle size and color encodes average value for percentage of PRC1 molecules found in condensates. For each condition 2.600 to >27.000 spots were analyzed.

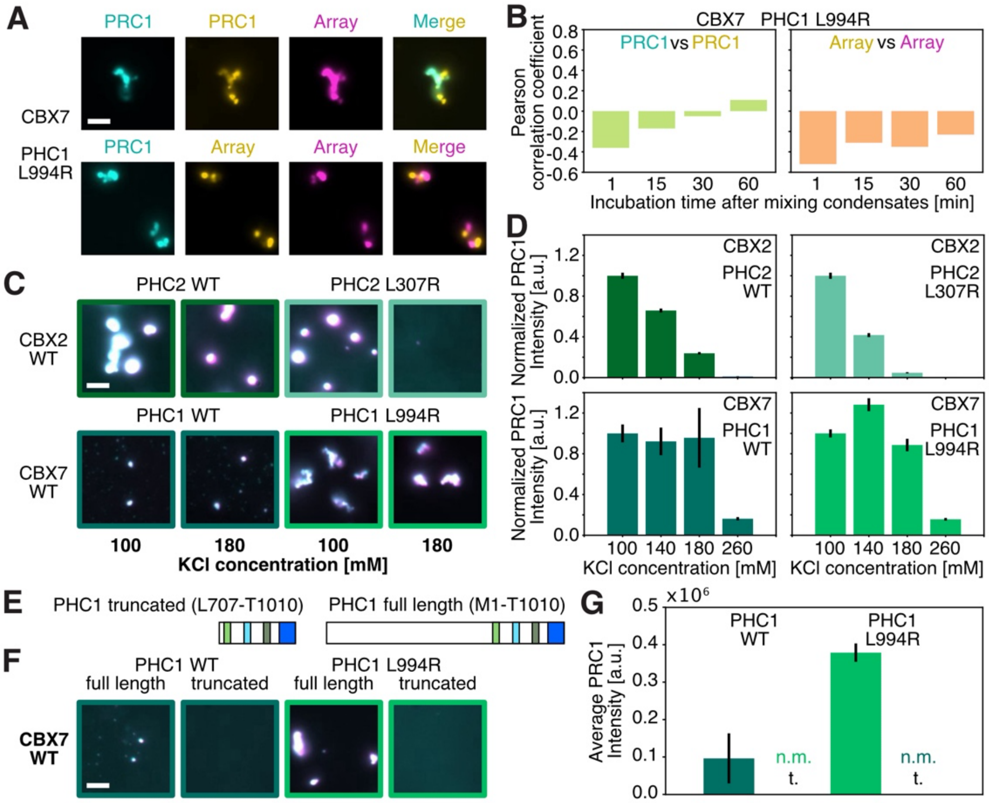


**Figure S4 | PHC1 alters condensate dynamics and stability.** (**A**) Representative micrographs of condensates with differently colored PRC1 molecules (top) and differently colored arrays (bottom) 30 minutes after mixing for PRC1 complexes containing CBX7 and PHC1 L994R. (**B**) Average Pearson correlation coefficients of differently colored PRC1 molecules (left) and nucleosomal arrays (right) for PRC1 complexes with CBX7 and PHC1 L994R. The nucleosome concentration was 30 nM and PRC1 concentration was 320 nM. For each condition >100 spots have been analyzed. (**C**) Example micrographs of nucleosomal arrays (magenta) and canonical PRC1 complexes (cyan) for different PRC1 compositions at various potassium chloride concentrations. Nucleosome concentration was 30 nM and PRC1 concentration was 640 nM. The scale bar is 5 µm. (**D**) Normalized PRC1 intensities of data as shown in **C**. Error bar is standard error of the mean. (**E**) Schematic of full length and truncated PHC1. (**F**) Example micrographs of 30 nM nucleosomal arrays (magenta) and 320 nM canonical PRC1 complexes (cyan) with different subunits incorporated. The scale bar is 5 µm. (**G**) Average PRC1 intensities for different PRC1 compositions as shown in **J**. Error bar is standard error of the mean. f.l. is full length PHC1 and t. is truncated PHC1. (**B, D, G**) For each condition 100 to >30.000 spots were analyzed. Data are representative of at least two technical repeats. N.m. is not measurable.

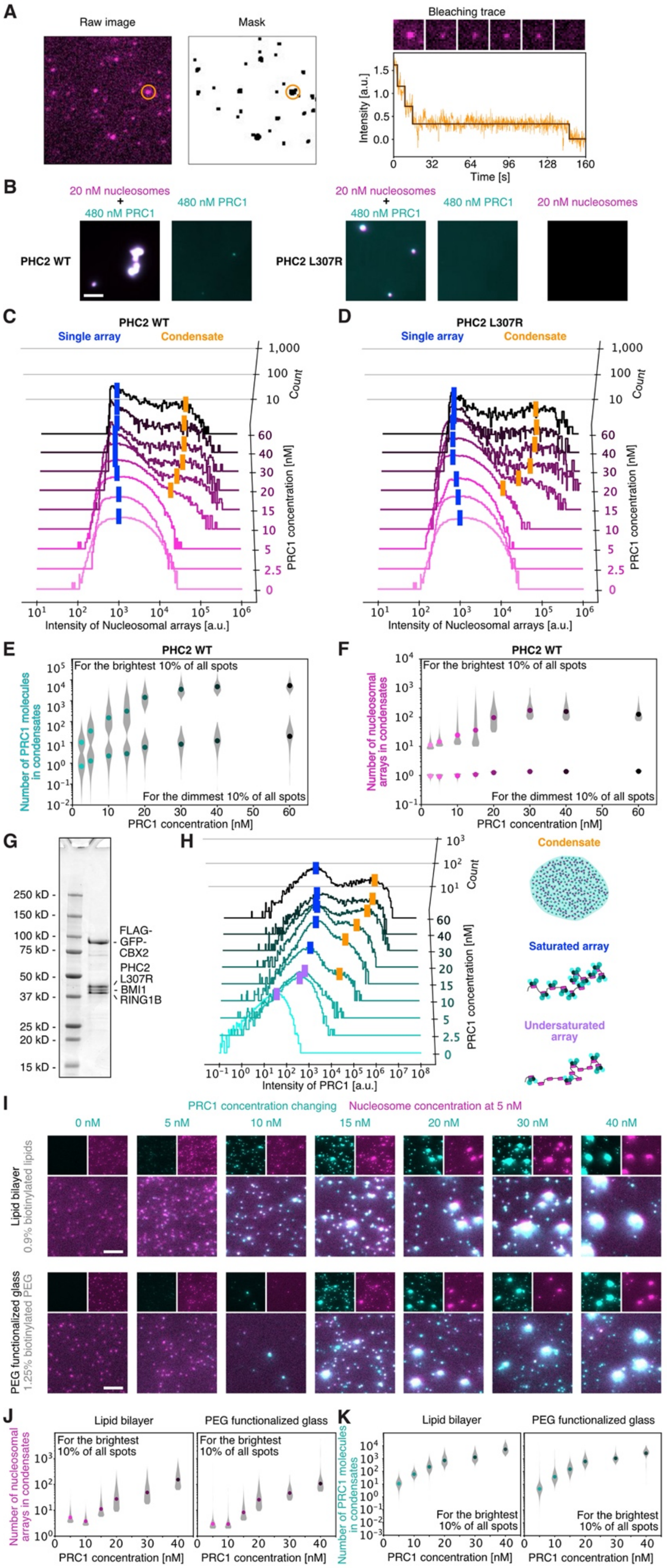
A
Raw image
Mask
Bleaching trace
Intensity [a.u.]
Time [s]
B
20 nM nucleosomes
+
480 nM PRC1
480 nM PRC1
20 nM nucleosomes
PHC2 WT
PHC2 L307R
C
PHC2 WT
Single array
Condensate
Count
PRC1 concentration [nM]
Intensity of Nucleosomal arrays [a.u.]
D
PHC2 L307R
E
PHC2 WT
For the brightest 10% of all spots
For the dimmest 10% of all spots
Number of PRC1 molecules in condensates
PRC1 concentration [nM]
F
Number of nucleosomal arrays in condensates
G
FLAG-GFP-CBX2
PHC2 L307R
BMI1
RING1B
H
Intensity of PRC1 [a.u.]
Condensate
Saturated array
Undersaturated array
I
PRC1 concentration changing
Nucleosome concentration at 5 nM
Lipid bilayer
0.9% biotinylated lipids
PEG functionalized glass
1.25% biotinylated PEG
J
Lipid bilayer
PEG functionalized glass
K

**Figure S5 | Additional experiments to validate TIRF assay to study PRC1 and nucleosomal array condensate formation.** (**A**) Example quantification of photobleaching analysis to estimate the number of nucleosomal arrays in condensates. (**B**) Example images of PRC1 complexes with PHC2 WT or PHC2 L307R alone, nucleosomal arrays alone, or when arrays and PRC1 are mixed. The images were taken after a 30-minute incubation at 30°C on an epifluorescence microscope. The scale bar is 5 µm. (**C, D**) Histograms of nucleosomal array intensities related to the PRC1 intensities shown in **Figure 7B** for (**C**) PHC2 WT and (**D**) PHC2 L307R. The blue rectangles indicate spots with single nucleosomal arrays while the orange rectangles show condensates. (**E, F**) Violin plots of the number of PRC1 molecules (**E**) and nucleosomal arrays (**F**) for the brightest and dimmest 10% of all spots from the distributions shown in **Figure 7B**. (**G**) 4-20% SDS-PAGE of purified canonical PRC1 complex with PHC2 L307R without HALO-tag. (**H**) Histograms of PRC1 intensities of PRC1 complexes with PHC2 L307R without the HALO-tag. The purple rectangles indicate spots with undersaturated nucleosomal arrays, the blue rectangles correspond to single, saturated nucleosomal arrays, and the oranges rectangles show condensates of nucleosomal arrays and PRC1. (**I**) Representative TIRF micrographs of PRC1 (cyan) and nucleosomal arrays (magenta) on biotinylated lipid bilayers (top) and on functionalized glass with biotinylated PEG (bottom). The scale bar is 5 µm. (**J, K**) Violin plots of the number of nucleosomal arrays (**J**) and PRC1 molecules (**K**) for the brightest and dimmest 10% of all spots on both surfaces. For each condition 2.500 to > 60.000 spots were analyzed. (**B-F, H**) Data are representative of at least three technical repeats. Note, **H-K** were performed with a PRC1 complex containing PHC2 L307R without HALO-tag.

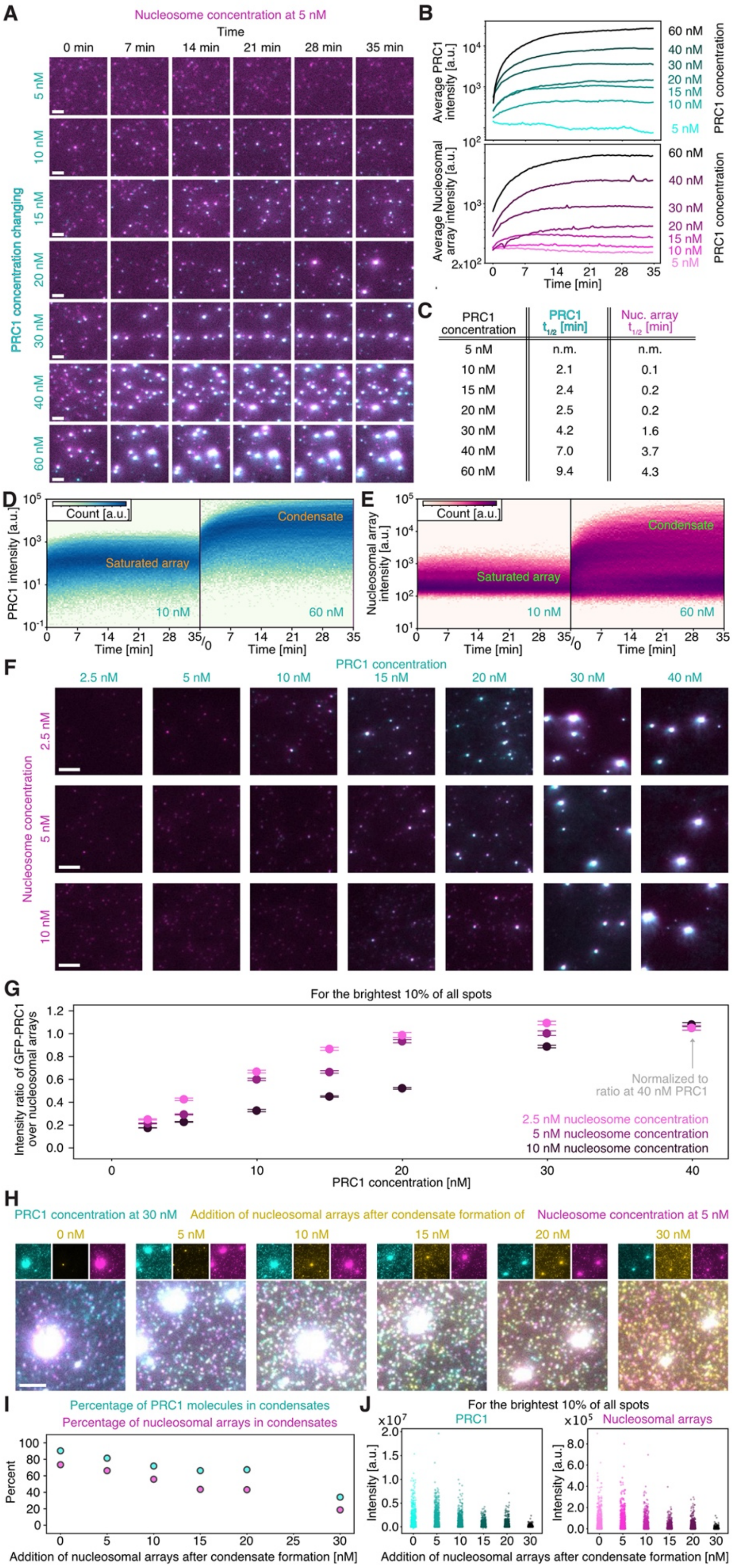


| PRC1 concentration | PRC1 $t_{1/2}$ [min] | Nuc. array $t_{1/2}$ [min] |
|---|---|---|
| 5 nM | n.m. | n.m. |
| 10 nM | 2.1 | 0.1 |
| 15 nM | 2.4 | 0.2 |
| 20 nM | 2.5 | 0.2 |
| 30 nM | 4.2 | 1.6 |
| 40 nM | 7.0 | 3.7 |
| 60 nM | 9.4 | 4.3 |

**Figure S6 | Switch-like behavior of PRC1-nucleosomal array condensate formation.** (**A**) Representative micrographs of condensate formation over time of PRC1 (cyan) and nucleosomal arrays (magenta). (**B**) Average intensity traces of all PRC1 (top) nucleosomal array (bottom) spots over time. (**C**) Quantification of half-time to reach maximum intensity for PRC1 and nucleosomal arrays of graphs shown in **B**. Data was fitted with a single exponential. (**D, E**) Heatmap of intensity distributions of (**D**) PRC1 and (**E**) nucleosomal arrays over time. (**F**) TIRF micrographs of PRC1 (cyan) and nucleosomal arrays (magenta) for various nucleosome and PRC1 concentrations. The scale bar is 5 µm. (**G**) Average intensity ratio of PRC1 over nucleosomal arrays normalized to the ratio at 40 nM PRC1. The error bar shows the standard error of the mean. (**H**) Example micrographs for conditions for which we added Cy3N labeled nucleosomal arrays to preformed condensates of ATTO-647N labeled nucleosomal arrays and mGFP-tagged PRC1. The scale bar is 5 µm. (**I**) Percentage of molecules in condensates of PRC1 (cyan) and nucleosomal arrays (magenta) when different amounts of additional arrays are added to preformed condensates. (**J**) Quantification of intensities of individual spots when different amounts of additional arrays are added to preformed condensates. (**B-E, G, I, J**) For each condition 2.000 to >100.000 spots were analyzed. Data are representative of at least two technical repeats. Note, all experiments for this figure were performed with a PRC1 complex containing PHC2 L307R without HALO-tag. This was necessary since we did not want to limit the condensation dynamics to the rate of TEV cleavage of the HALO-tag of PHC2.

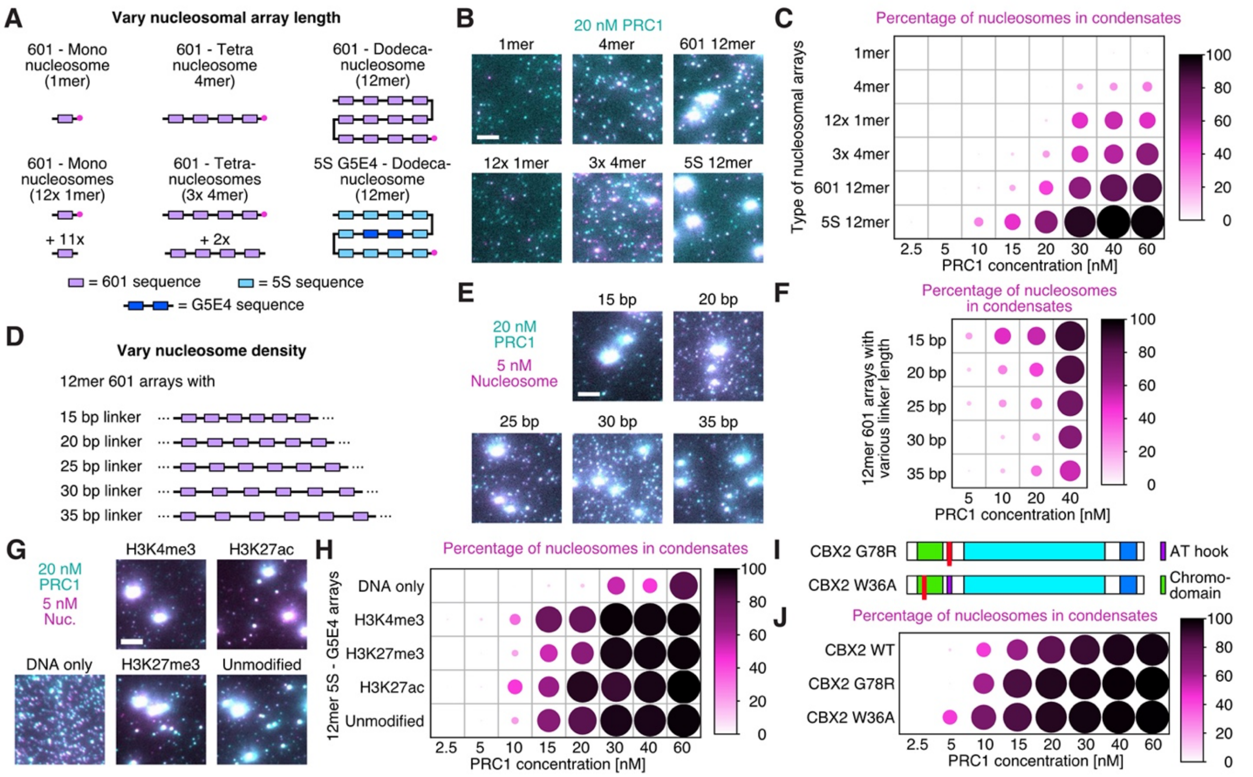


**Figure S7 | Influence of various nucleosomal array parameters on condensate formation with PRC1.** (**A, D**) Schematics of nucleosomal arrays of various lengths (**A**) and with different linker DNA lengths between nucleosomes (**D**). (**B, E**) Micrographs of PRC1 (cyan) and nucleosomal arrays (magenta) of various lengths (**B**) and different linker lengths (**E**). The scale bar is 5 µm. In **B** the 1mer, 4mer, 601 12mer, and 5S 12mer all have the same DNA concentration while for 12x 1mer, 3x 4mer, 601 12mer, and 5S 12mer the nucleosome concentration is the same and at 5 nM. (**C, F**) Percentage of nucleosomal arrays in condensates. (**G**) Micrographs of PRC1 (cyan) and nucleosomal arrays (magenta) with various histone modifications or for DNA alone. The scale bar is 5 µm. (**H**) Percentage of nucleosomal arrays in condensates for nucleosomal arrays with various histone modifications or for DNA alone. (**C, F, H**) The nucleosome concentration was 5 nM. Circle size and color encode average value for percentage of nucleosomal arrays found in condensates from three repeats. For each condition 500 to >100.000 spots were analyzed. (**I**) Schematics of CBX2 subunit with mutations in the chromodomain (W36A) and AT hook (G78R). (**J**) Percentage of nucleosomal arrays in condensates for full canonical PRC1 complex with different CBX2 subunit mutations. The nucleosome concentration was 5 nM. Circle size and color encode average value for percentage of nucleosomal arrays found in condensates. For each condition 2.600 to >27.000 spots were analyzed. (**A-J**) Note, all experiments for this figure were performed with a PRC1 complex containing PHC2 L307R without HALO-tag.

### Legends for Supplementary Movies

**Movie S1**
The movie shows PRC1 complexes with CBX2 and PHC2 WT (cyan) and nucleosomal array (magenta) before and after bleaching as shown in **Figure 2F**. Time is shown as minutes:seconds. The scale bar is 2 μm.

**Movie S2**
The movie shows PRC1 complexes with CBX2 and PHC2 L307R (cyan) and nucleosomal array (magenta) before and after bleaching as shown in **Figure 2F**. Time is shown as minutes:seconds. The scale bar is 2 μm.

**Movie S3**
The movie shows PRC1 complexes with CBX2 and PHC2 WT (cyan) and nucleosomal array (magenta) before and after bleaching as shown in **Figure S2I**. Time is shown as minutes:seconds. The scale bar is 2 μm.

**Movie S4**
The movie shows PRC1 complexes with CBX2 and PHC2 L307R (cyan) and nucleosomal array (magenta) before and after bleaching as shown in **Figure S2J**. Time is shown as minutes:seconds. The scale bar is 2 μm.

**Movie S5**
The movie shows examples of condensate formation over time of PRC1 (cyan) and nucleosomal arrays (magenta). Time is shown as minutes:seconds. The scale bar is 10 μm.

### Supplementary References


1. Kim, J.J. Context-specific Polycomb mechanisms in development. 16.
2. Blackledge, N.P., and Klose, R.J. (2021). The molecular principles of gene regulation by Polycomb repressive complexes. Nat. Rev. Mol. Cell Biol. *22*, 815–833. 10.1038/s41580-021-00398-y.
3. Flaus, A. (2011). Principles and practice of nucleosome positioning in vitro. Front. Life Sci. *5*, 5–27. 10.1080/21553769.2012.702667.
4. Utley, R.T., Ikeda, K., Grant, P.A., Côté, J., Steger, D.J., Eberharter, A., John, S., and Workman, J.L. (1998). Transcriptional activators direct histone acetyltransferase complexes to nucleosomes. Nature *394*, 498–502. 10.1038/28886.
5. Plys, A.J., Davis, C.P., Kim, J., Rizki, G., Keenen, M.M., Marr, S.K., and Kingston, R.E. (2019). Phase separation of Polycomb-repressive complex 1 is governed by a charged disordered region of CBX2. Genes Dev. *33*, 799–813. 10.1101/gad.326488.119.
6. Lau, M.S., Schwartz, M.G., Kundu, S., Savol, A.J., Wang, P.I., Marr, S.K., Grau, D.J., Schorderet, P., Sadreyev, R.I., Tabin, C.J., et al. (2017). Mutation of a nucleosome compaction region disrupts Polycomb-mediated axial patterning. Science *355*, 1081–1084. 10.1126/science.aah5403.
7. Grau, D.J., Chapman, B.A., Garlick, J.D., Borowsky, M., Francis, N.J., and Kingston, R.E. (2011). Compaction of chromatin by diverse Polycomb group proteins requires localized regions of high charge. Genes Dev. *25*, 2210–2221. 10.1101/gad.17288211.
